# Success versus failure in cognitive control: meta-analytic evidence from neuroimaging studies on error processing

**DOI:** 10.1101/2023.05.10.540136

**Authors:** Edna C. Cieslik, Markus Ullsperger, Martin Gell, Simon B. Eickhoff, Robert Langner

**Author notes:** Correspondence should be addressed to: Edna C. Cieslik, Institute of Neuroscience and Medicine (INM-7), Research Centre Jülich, D-52425 Jülich, Germany.

## Abstract

Brain mechanisms of error processing have often been investigated using response interference tasks and focusing on the posterior medial frontal cortex, which is also implicated in resolving response conflict in general. Thereby, the role other brain regions may play has remained undervalued. Here, activation likelihood estimation meta-analyses were used to synthesize the neuroimaging literature on brain activity related to committing errors versus responding successfully in interference tasks and to test for commonalities and differences. The salience network and the temporoparietal junction were commonly recruited irrespective of whether responses were correct or incorrect, pointing towards a general involvement in coping with situations that call for increased cognitive control. The dorsal posterior cingulate cortex, posterior thalamus, and left superior frontal gyrus showed error-specific convergence, which underscores their consistent involvement when performance goals are not met. In contrast, successful responding revealed stronger convergence in the dorsal attention network and lateral prefrontal regions. Underrecruiting these regions in error trials may reflect failures in activating the task-appropriate stimulus-response contingencies necessary for successful response execution.

## 1. Introduction

Flexible, adaptive behaviour requires monitoring the appropriateness and outcome of our actions, including the detection of erroneous responses (i.e. error monitoring) and adjustment of the following behaviour. When errors inevitably occur, they aid optimization by indicating where adjustments to behaviour are necessary (Ullsperger et al., 2014a). Investigating error-related processes can thus offer a window into how performance is monitored in the brain. Conversely, failures of such a system resulting in increased error rates have been described as impulsivity (Moeller et al. 2001; Castellanos-Ryan et al., 2011), a behavioural tendency related to psychiatric conditions such as attention deficit hyperactivity disorder (ADHD) and linked with maladaptive behaviours such as smoking (Sharma et al., 2014).

According to Reasońs influential Generic error-modelling system (GEMS) (Reason, 1990), three different types of errors can be observed in humans: (i) skill-based slips and lapses, (ii) rule-based mistakes, and (iii) knowledge-based mistakes. Skill-based errors occur when the intention is correct but carrying out the required action fails (cf. Norman, 1981). In contrast, mistakes are defined as actions that run according to plan, but the plan itself is inadequate to achieve a desired outcome. Mistakes are therefore often interpreted as problem solving failures.

In this study, we focus on skill-based errors, which in the psychological literature have often been studied using tasks in which two response tendencies collide. In such tasks, typically, a predominant but inadequate response tendency competes and interferes with a non-dominant but adequate (instructed) response. The conflict, in turn, can lead to the execution of an inadequate response (i.e. a commission error) on a substantial number of trials. Two of the most commonly employed tasks to study overriding a predominant motor response are the go/no-go (Donders, 1969) and stop-signal tasks (Logan et al., 1997), which require to withhold or cancel the predominant go-response. In another set of tasks, such as the Stroop, flanker, Simon, or stimulus–response compatibility (SRC) tasks, a given stimulus property interferes with relevant stimulus and/or response information, impeding responses to the relevant information. Usually, responses are faster and more accurate if stimulus (S) and response (R) features correspond as compared to when they do not (e.g. Fitts and Seeger, 1953, Hallett, 1978; Eimer et al., 1995; Cieslik et al., 2010). Successful behaviour in all these tasks is therefore assumed to require the suppression of the automatically activated predominant response tendency, while concurrently implementing the context-appropriate one. This can be either not responding at all or activating the schema for the incongruent alternative (cf. Cieslik et al., 2015). If activation of the context-appropriate response fails or is not achieved fast enough, commission errors occur.

The neural correlates of erroneous responding and subsequent behavioural adjustments have been extensively studied using electroencephalography (EEG). Here, two event-related components are commonly observed after commission errors. First, the error-related negativity (ERN) (Gehring et al., 1993) or error negativity (Ne) (Falkenstein et al., 1990) can be observed, with a negative peak at 50-100 ms after the initiation of the error response. This component is most likely generated within the posterior medial frontal cortex (pMFC) (Dehaene et al., 1994; Debener et al., 2005; Ridderinkhof et al., 2004; Ullsperger and von Cramon, 2001; Fu et al., 2022), including the anterior midcingulate cortex (aMCC) and extending into the pre-supplementary motor area (preSMA). Functionally, the ERN has been argued to reflect processing a mismatch between correct and actual responses (Scheffers and Coles, 2000), a post-response conflict (Botvinick et al., 2001; Yeung et al., 2004), or a reinforcement learning signal (Holroyd and Coles, 2002). All of these slightly different accounts share the idea that the ERN is based on detecting that the current response differs from the required (correct) one. In line with that, a recent study showed the ERN to depend on the representation of the correct response to emerge (Di Gregorio et al., 2018). This performance monitoring response is then thought to be conveyed to other regions, such as lateral PFC (MacDonald et al., 2000), for implementing adaptation processes to avoid the same error in the future.

These post-error adjustments show in different behavioural phenomena (for a review, see Danielmeier and Ullsperger, 2011): (i) post-error slowing (PES), i.e. longer RT in trials after committing an error than in trials after correct responding (Rabbitt, 1966; Laming, 1968); (ii) post-error reduction of interference (PERI), i.e. smaller interference costs in trials following errors than in trials following correct responses (King et al., 2010; Ridderinkhof, 2002); and (iii) post-error improvement in accuracy (PIA), i.e. higher accuracy in trials after errors than after correct responses (Danielmeier et al., 2011, Laming, 1968).

Following the ERN, a second, positive deflection, the error positivity (Pe) has been repeatedly observed, peaking around 200-500 ms after committing the error (Falkenstein et al., 1990). It has been argued that the Pe reflects a neural index of error awareness (O’Connell et al., 2007; Hughes and Yeung, 2011; Steinhauser and Yeung, 2010; Ullsperger et al., 2014b; Kirschner et al., 2021). The localization of the neural generators of this component are, however, less clear, with some evidence pointing to multiple sources, including rostral pMFC (van Veen and Carter, 2002), caudal pMFC (Herrmann et al., 2004), the parietal cortex (Herrmann et al., 2004), and also the dorsolateral prefrontal cortex (cf. Masina et al., 2019).

Potential causes for erroneous behaviour have been related to attentional lapses or control failures, reflected, for example, in increased activity of the default-mode network (DMN) in trials preceding an error (e.g. Li et al., 2007; Weissman et al., 2006; Su et al., 2020). There is, however, also evidence from event-related EEG that error-preceding activity may reflect an adaptive adjustment of cognitive control (Eichele et al., 2008, 2010). This, paradoxically, may itself contribute to the generation of error-prone states because a strong reduction in cognitive control after presentation of several compatible trials may then lead to an error on a following incompatible trial (cf. Steinhauser et al., 2012). To summarise, there is evidence, in particular from EEG studies, about error-preceding processes as well as the neuronal generators of error-related activity. While electrophysiological recordings provide very high temporal resolution that help to better disentangle different processes on a temporal scale, spatial resolution is relatively coarse, unless combined with other approaches, such as functional magnetic resonance imaging (fMRI). In particular, the neural origins of Pe are not well understood, as multiple brain regions may contribute to this signal.

fMRI studies as well as one previous neuroimaging meta-analysis on error processing, which included only a relatively small number of experiments, have strongly focussed on the pMFC (Carter et al., 1998; Garavan et al., 2003; Critchley et al., 2005; Sharp et al., 2010; Ullsperger and von Cramon, 2001; Klein et al., 2007) and the anterior insula (aI) (Magno et al., 2006; Ullsperger et al., 2010), often glossing over the roles other brain regions with significant error-related activity may play. Moreover, a systematic comparison to the network involved when the task at hand is performed correctly is missing. This comparison, however, is crucial for a better understanding of which regions are specific to error processing and which regions are more generally involved in interference resolution. With the present study, we aimed to fill that gap by isolating and functionally characterising brain regions that are consistently linked to the detection of errors and feeding this information on to other brain regions associated with adaptation processes in subsequent trials. Moreover, as some authors have argued that error-related activity in the pMFC reflects the detection of conflicting response plans rather than an error-specific response (e.g. Botvinick et al., 2001; Yeung et al., 2004), we furthermore compared the network for error processing with the network consistently involved in performing the above-mentioned interference tasks successfully. To this end, coordinate-based activation likelihood estimation (ALE) meta-analyses (Eickhoff et al., 2012; Eickhoff et al., 2009; Turkeltaub et al., 2002; Turkeltaub et al., 2012) were used to integrate results from a diverse range of neuroimaging studies investigating (i) *commission errors* and (ii) *successful interference control* in tasks requiring speeded responding, in particular the go/no-go, stop-signal, Stroop, flanker, manual SRC, antisaccade, and Simon tasks. ALE meta-analysis allows testing for brain regions consistently involved in a mental function across different studies, independent of specific task variants (cf. Müller et al., 2018). Due to the rather coarse temporal resolution of fMRI, activity during commission error trials does most likely not only reflect error detection per se but also processes signalling the need for subsequent cognitive adjustments to other brain regions and potentially also the implementation of these processes.

Performing a minimum conjunction analysis across the meta-analyses on *commission errors* and *successful interference control* allowed us to reveal brain regions consistently activated whenever the task context requires inhibiting the predominant response and concurrently activating the appropriate task goal for initiating the adequate behaviour, independent of the behavioural outcome. In contrast, meta-analytic contrast analyses would reveal those brain regions specifically involved during commission errors or successful interference resolution, respectively. Using this approach, we tried to improve our understanding of which regions are more specifically involved in error-related processes and which brain regions may be crucial for activating the correct, non-automatic behavioural alternative.

## 2. Methods

### 2.1 Paradigms included

For our meta-analysis, neuroimaging results on the neural correlates of correct as well as erroneous responses in seven different tasks investigating interference control were included, namely the go/no-go, stop-signal (Logan, 1997), Stroop (Stroop, 1935), flanker (Eriksen, 1974), Simon (Simon, 1990), antisaccade (Hallett, 1978) and manual SRC task. Importantly, all of these tasks require the implementation of a non-dominant, context-dependent behaviour against a competing behavioural alternative. In the go/no-go and stop-signal tasks, a dominant tendency for a speeded response (versus inhibiting an overt response) is commonly achieved by presenting go (or no-stopping) trials with a higher frequency than inhibition (i.e., no-go or stop) trials (cf. Eagle et al., 2008; Schachar et al., 2007). In the Stroop, flanker, Simon, antisaccade and manual SRC tasks, a given stimulus dimension interferes with relevant stimulus and/or response information, impeding responses to the relevant dimension.

### 2.2. Selection criteria for the experiments included in the meta-analyses

The present meta-analyses were based on the database of a previous meta-analytic study on interference control (Cieslik et al., 2015). Publications within this database were screened for experiments focussing on commission errors as well as successful interference control. Additionally, a literature search using PubMed (www.pubmed.com) and Google Scholar https://scolar.google.de) was performed to obtain additional relevant neuroimaging experiments. Further experiments were detected by reference tracing in the retrieved papers and in review articles.

We only included results from whole-brain group analyses based on data from healthy adults (age ≥ 18 years) reported as coordinates in a standard reference space (Talairach/Tournoux or Montreal Neurological Institute (MNI)). Reported coordinates of experiments using FSL or SPM were treated as MNI coordinates as this is the standard space in these softwares. Only in cases when authors explicitely reported a transformation of MNI in TAL space, the coordinates were considered to be in TAL space when using FSL or SPM. Otherwise the coordinate space was treated as MNI (cf. Müller et al., 2018).

Results of region-of-interest (ROI) analyses were excluded. When clinical studies separately reported results from a healthy control group (i.e., excluding data from patients), these results were included. Data from conditions with pharmacological or non-invasive manipulations such as Transcranial Magnetic Stimulation were excluded. Importantly, if the same participants or a subsample of the same participants was used in several studies, only one of the studies was included (e.g. Ray Li et al., 2006a,b) to control for possible sample-specific effects. If, however, one publication separately reported results from different participant samples, such as young and old participants (e.g. Korsch et al., 2014), the respective contrasts were handled as two independent experiments in the meta-analysis. Further details on inclusion and exclusion criteria are provided in the checklist in the supplementary material.

For the meta-analyses on error processing, we included contrasts reporting commission errors > successful interference resolution (e.g., commission error > successful incongruent trials in the Stroop task, or commission error > successful stop trials in the stop-signal task) as well as contrasts reporting commission errors > a control condition (e.g., commission errors > congruent, commission errors > go, or commission errors > low-level baseline condition). For the meta-analysis on successful interference resolution, we included experiments contrasting correct interference resolution > no interference (i.e., correct stop > go, or correct incongruent > congruent). Experimental effects compared with a resting baseline as well as the reverse contrasts (i.e., “deactivations”) were not considered for inclusion. However, in some of the go/no-go experiments, go trials were not modelled explicitly, but included in an “active” baseline. We included these experiments as we considered the “baseline” as an active control condition in these experiments. Importantly, we here focused on commission errors as omission errors are very rare in these paradigms and are usually modelled as a regressor of no interest at the single-subject level only.

Based on the criteria defined above, 43 experiments on commission errors and 142 experiments on successful interference control (i.e., response conflict processing) were included in the meta-analyses. Experiments from the same publication were pooled to account for sample overlap (Turkeltaub et al., 2012). To account for differences in coordinate space (MNI vs. Talairach space), coordinates reported in Talairach space were converted into MNI coordinates using a linear transformation (Lancaster et al., 2007). See table S1 and S2 for a full overview of all included studies and further information such as number of participants or which contrast was used in the respective study.

We performed two meta-analyses: (1) a meta-analysis on *commission errors* including experiments contrasting commission errors versus successful interference control or versus control condition or baseline (n = 43 experiments), and (2) a meta-analysis on *successful interference control* including experiments contrasting correct interference responses > no-interference responses (n= 142 experiments). Further, all resulting clusters of convergence were analyzed with regard to which experiments contributed to the specific effect. An overview of the contributing experiments is provided in the supplementary material.

To identify regions consistently involved in both processes, a minimum conjunction analysis was performed, while a meta-analytic contrast analysis between these two meta-analyses was calculated to look for consistent differences between conditions. A full overview of the workflow is provided in the flowchart in the supplementary material (Fig. S1).

To account for the difference in sample size between the two analyses, we replicated the approach with a reduced, matched sample, only including experiments from publications that reported both contrasts, resulting in 30 experiments for commission error and 30 experiments for correct interference resolution, respectively. These results are provided in the supplementary material.

Additionally, we performed two control meta-analyses on commission errors separating between the contrasting condition: (1) commission errors > successful interference control (n = 24), and (2) commission errors versus no-interference responses or low-level baseline (n = 22). Further, we conducted two additional meta-analyses separating between paradigm classes: (1) errors during classic response inhibition tasks (i.e. go/no-go and stop signal), and (2) errors during the classic response interference tasks (Stroop, flanker, Simon, antisaccade and manual SRC task). Importantly, for the latter two meta-analyses, we did not only include experiments contrasting commission errors versus successful interference control/no-interference/low-level baseline but also experiments that did not specifically model commission errors but included commission as well omission errors in the regressor of interest. The reason behind this approach was the rather low number of experiments specifically modelling only commission errors in the Stroop, flanker, Simon or antisaccade tasks. As mentioned before, in such interference tasks, most errors occur in the incongruent condition, while omission errors are rarely observed. Hence, this should not have an impact on the results.

All results are made available in the ANIMA database (https://https://anima.fz-juelich.de) (Reid et al., 2016).

### 2.3 Activation Likelihood Estimation

All meta-analyses were performed using the revised ALE algorithm for coordinate-based meta-analysis (Eickhoff et al., 2012; Eickhoff et al., 2009; Laird et al., 2009a; Laird et al., 2009b; Turkeltaub et al., 2002) according to the standard procedures of our institute (see, e.g., Langner and Eickhoff, 2013; Heckner et al., 2021; Müller et al., 2018 for a more detailed description). ALE aims to identify areas showing convergence of reported coordinates across experiments that is higher than expected under the assumption of random spatial associations. Therefore, the probabilities of all foci reported in a given experiment were combined for each voxel, resulting in a modelled activation (MA) map (Turkeltaub et al., 2012). Taking the union across these MA maps yielded voxel-wise ALE scores describing the convergence of results across experiments at each particular location of the brain. To distinguish ‘true’ convergence across studies from random convergence (i.e., noise), ALE scores were compared to an empirical null-distribution reflecting a random spatial association between experiments. The p-value of the “true” ALE was then given by the proportion of equal or higher values obtained under the null-distribution. The resulting non-parametric p-values for each meta-analysis were thresholded at a cluster-level corrected threshold of p < 0.05 (cluster-forming threshold at voxel level: p < 0.001, uncorrected) and transformed into z-scores for display.

To identify voxels showing a significant effect in both separate analyses, the conservative minimum conjunction statistic (Nichols et al., 2005) was used. Further, an additional extent threshold of k ≥ 25 was applied to exclude smaller clusters that most likely reflect incidental overlap between significant effects in both meta-analyses.

Meta-analytic contrast analyses were performed to reveal voxels showing consistently stronger convergence in one meta-analysis than the other (for a more detailed description of contrast analyses, see Langner and Eickhoff, 2013; Rottschy et al., 2012). Differences in ALE scores between two conditions were tested against a voxel-wise null-distribution of label exchangeability and thresholded at a posterior probability of p > 95% for true differences. Surviving voxels were inclusively masked by the respective main effect, i.e., the significant effect of the ALE analysis for the minuend, and an additional cluster extent threshold of k ≥ 25 voxels was applied.

All results were anatomically labelled by reference to probabilistic cytoarchitectonic maps of the human brain using the SPM Anatomy Toolbox version 3.0 (Eickhoff et al., 2007; Eickhoff et al., 2005).

## 3. Results

### 3.1 Meta-analysis on commission errors

Across 43 experiments reporting brain activity specifically related to committing errors during interference resolution, significant convergence was observed in bilateral anterior insula (aI), left superior frontal cortex, posterior medial frontal cortex (pMFC) including aMCC and preSMA, bilateral anterior temporo-parietal junction (TPJ), dorsal posterior cingulate cortex (PCC) as well as right V1 (hOc1). Subcortical convergence of activity was found in the posterior part of the thalamus (Fig. 1, Table 1).

**Figure 1:**
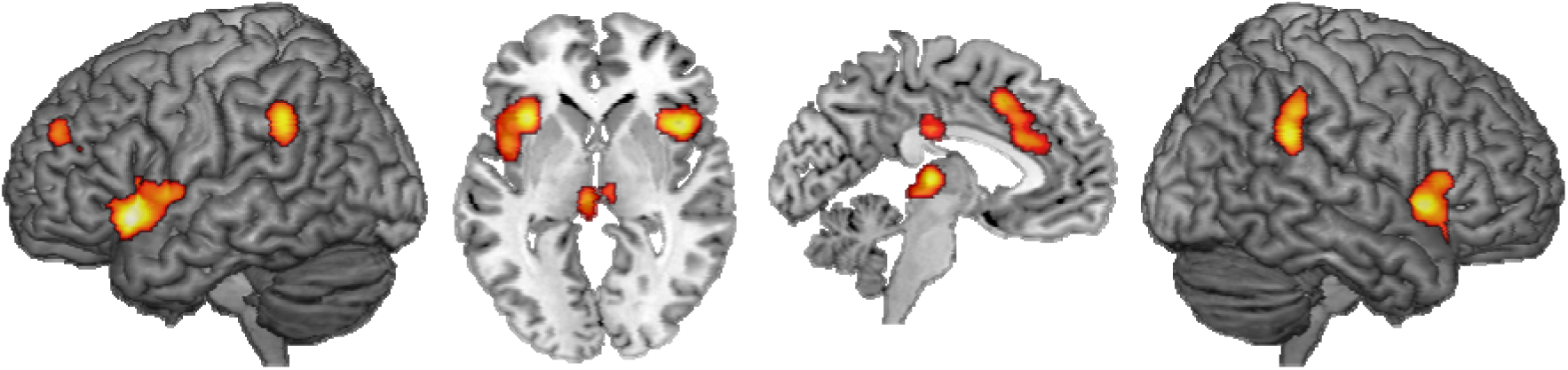
Brain regions showing significant convergence of activity for committing an error in high-conflict trials. Results are cluster-level p < .05 family-wise error-corrected for multiple comparisons, with a cluster forming threshold of p < .001 at the voxel level.

**Table 1:** Regions showing consistent activity across commission errors on a cluster-level corrected threshold of p<0.05 (with a cluster forming threshold of p<0.001).

| Cluster/Macroanatomical Structure | x | y | z | Histological assignment | z-score |
| --- | --- | --- | --- | --- | --- |
| **Cluster 1 (1548 voxels)** |  |  |  |  |  |
| Posterior medial frontal cortex (pMFC) | 4 | 20 | 40 |  | 7.18 |
| pMFC | 4 | 20 | 46 |  | 6.97 |
| pMFC | 6 | 32 | 28 |  | 6.53 |
| pMFC | 0 | 24 | 26 |  | 6.07 |
| pMFC | 2 | 14 | 60 |  | 3.84 |
| pMFC | 8 | 36 | 12 |  | 3.59 |
| **Cluster 2 (1189 voxels)** |  |  |  |  |  |
| Left anterior insula (aI) | -38 | 20 | -8 |  | 7.62 |
| Left medial insula | -44 | 4 | 4 |  | 5.04 |
| **Cluster 3 (761 voxels)** |  |  |  |  |  |
| Right aI | 42 | 14 | -2 |  | 6.82 |
| Right aI | 34 | 20 | -14 |  | 5.29 |
| Right aI | 32 | 22 | -12 |  | 5.29 |
| Right inferior frontal cortex | 50 | 22 | 8 | Area 45 | 4.85 |
| **Cluster 4 (387 voxels)** |  |  |  |  |  |
| Right inferior parietal cortex (IPC) | 60 | -42 | 28 | Area PFm | 6.29 |
| Right IPC | 54 | -40 | 38 | Area PFm | 5.28 |
| **Cluster 5 (293 voxels)** |  |  |  |  |  |
| Left posterior thalamus | -4 | -22 | 4 |  | 6.35 |
| Right posterior thalamus | 8 | -18 | 4 |  | 4.34 |
| Right medial thalamus | 12 | -10 | 4 |  | 3.57 |
| Right posterior thalamus | 8 | -24 | -2 |  | 3.39 |
| **Cluster 6 (220 voxels)** |  |  |  |  |  |
| Left IPC | -60 | -44 | 36 | Area PF | 5.92 |
| **Cluster 7 (160 voxels)** |  |  |  |  |  |
| Left superior frontal gyrus (SFG) | -28 | 50 | 28 |  | 4.43 |
| Left SFG | -24 | 46 | 26 |  | 4.15 |
| Left SFG | -26 | 44 | 24 |  | 4.08 |
| Left SFG | -28 | 42 | 22 |  | 4.06 |
| **Cluster 8 (150 voxels)** |  |  |  |  |  |
| Posterior cingulate cortex (PCC) | 0 | -20 | 30 |  | 5.41 |
| **Cluster 9 (93 voxels)** |  |  |  |  |  |
| Primary visual cortex | 14 | -64 | 6 | Area hOc1 | 4.93 |

### 3.2 Meta-analysis on successful interference resolution

The meta-analysis across all 142 experiments investigating successful interference resolution revealed significant convergence of activity in a broad bilateral fronto-parietal network comprising aI and adjacent inferior frontal gyrus; lateral prefrontal cortex (lPFC) covering inferior frontal junction (IFJ), parts of dorsolateral prefrontal cortex (DLPFC), and dorsal premotor cortex (dPMC); aMCC/preSMA; and posterior parietal cortex spanning from superior parietal lobule (SPL) to intraparietal sulcus (IPS) and inferior parietal cortex (IPC)/TPJ. Moreover, significant convergence was found in the left inferior occipital cortex (IOC) and, subcortically, caudate and thalamus (Fig. 2, Table 2).

**Figure 2:**
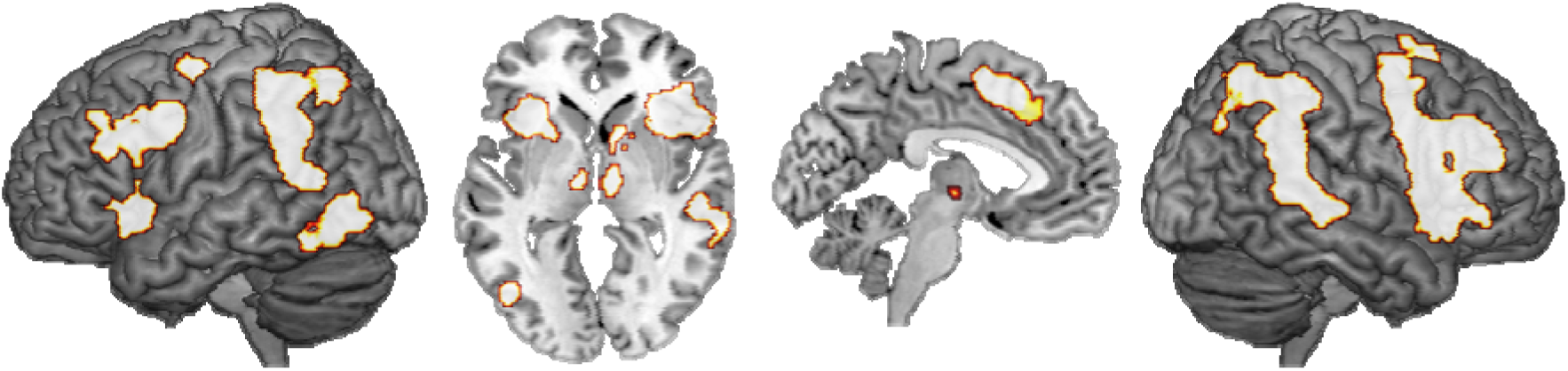
Brain regions showing significant convergence of activity for successful high-conflict trials. Results are cluster-level p < .05 family-wise error-corrected for multiple comparisons, with a cluster forming threshold of p < .001 at the voxel level.

**Table 2:**
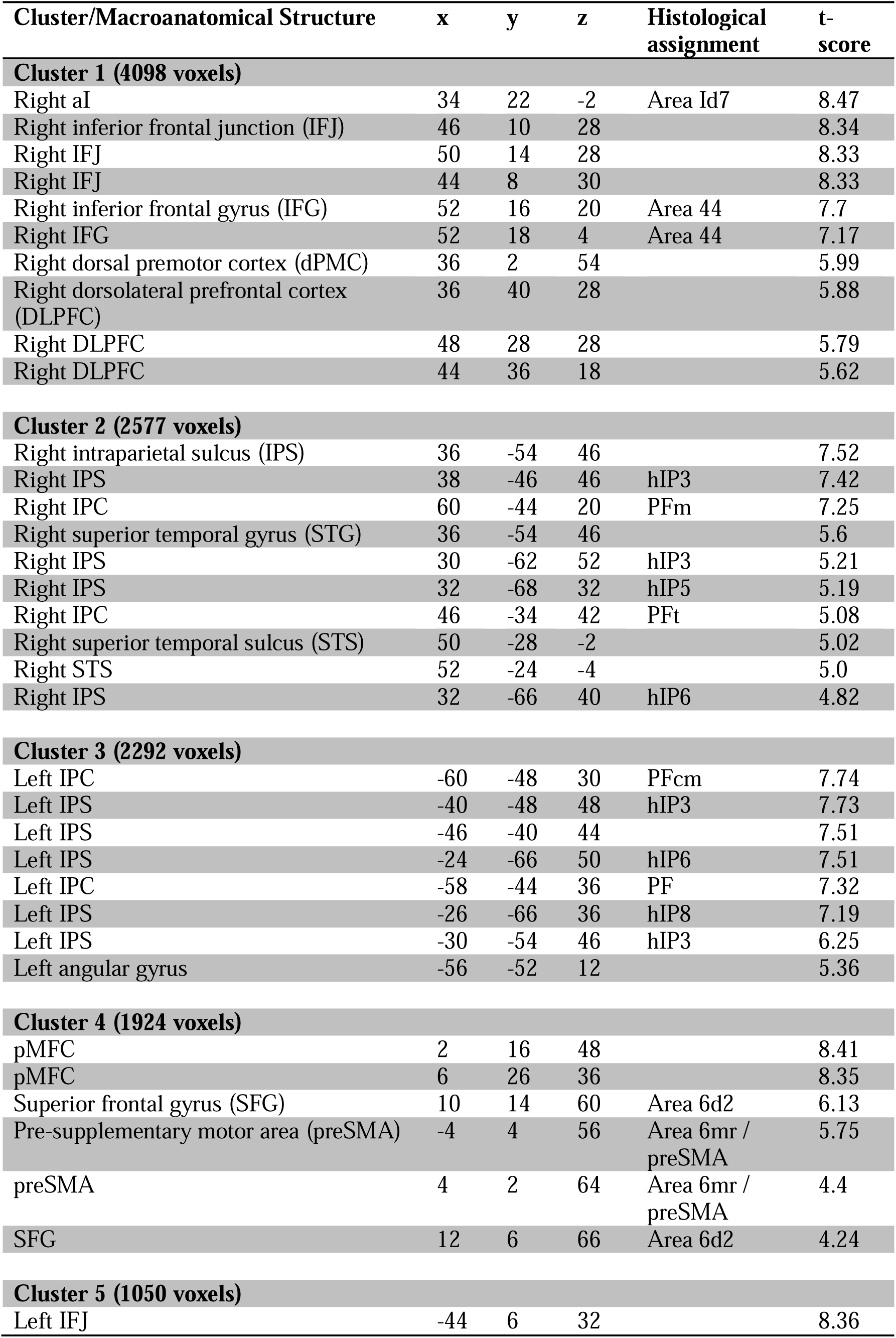
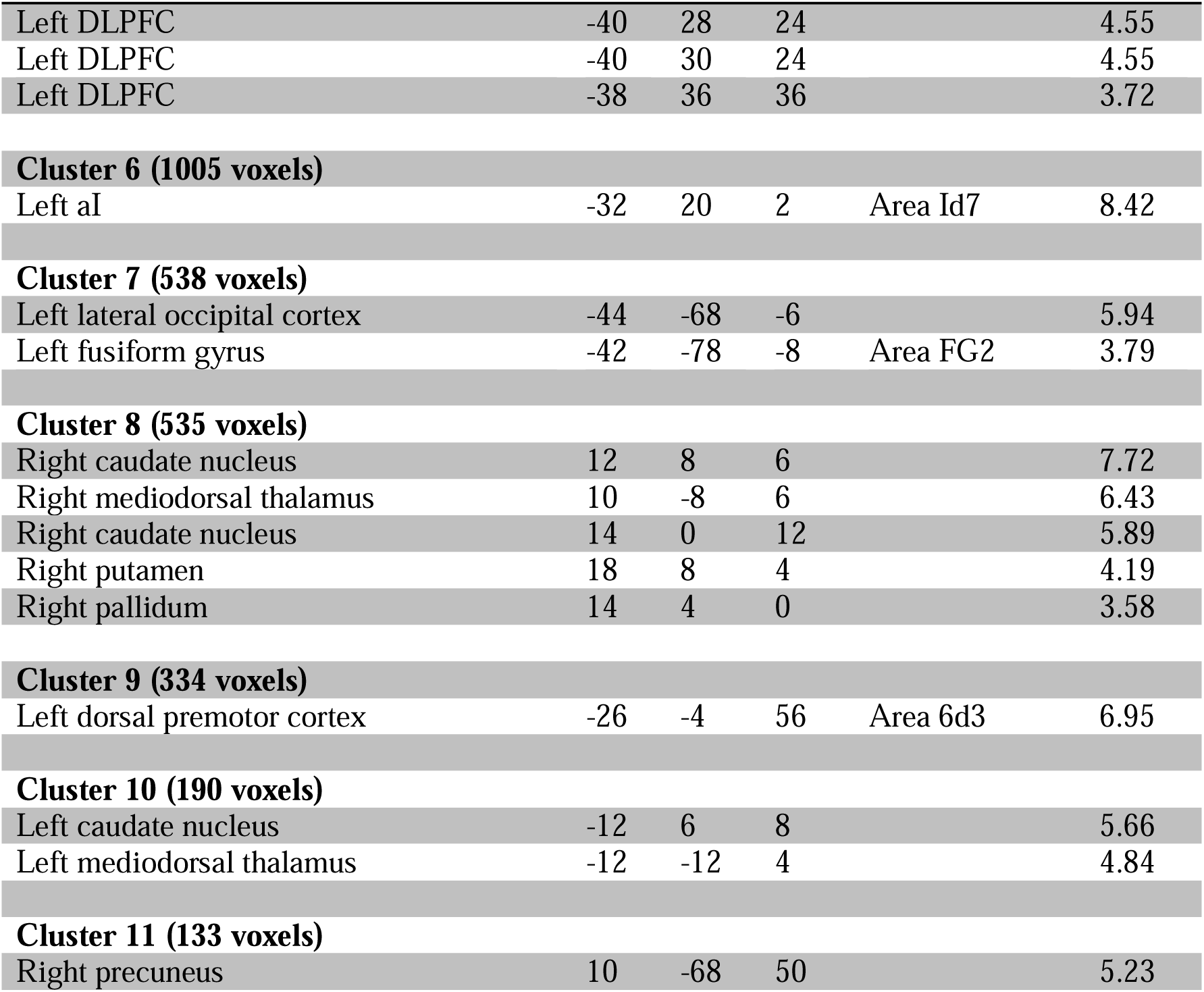
Regions showing consistent activity for correct interference resolution on a cluster-level corrected threshold of p<0.05 (with a cluster forming threshold of p<0.001)

| Cluster/Macroanatomical Structure | x | y | z | Histological assignment | t-score |
| --- | --- | --- | --- | --- | --- |
| **Cluster 1 (4098 voxels)** |  |  |  |  |  |
| Right aI | 34 | 22 | -2 | Area Id7 | 8.47 |
| Right inferior frontal junction (IFJ) | 46 | 10 | 28 |  | 8.34 |
| Right IFJ | 50 | 14 | 28 |  | 8.33 |
| Right IFJ | 44 | 8 | 30 |  | 8.33 |
| Right inferior frontal gyrus (IFG) | 52 | 16 | 20 | Area 44 | 7.7 |
| Right IFG | 52 | 18 | 4 | Area 44 | 7.17 |
| Right dorsal premotor cortex (dPMC) | 36 | 2 | 54 |  | 5.99 |
| Right dorsolateral prefrontal cortex (DLPFC) | 36 | 40 | 28 |  | 5.88 |
| Right DLPFC | 48 | 28 | 28 |  | 5.79 |
| Right DLPFC | 44 | 36 | 18 |  | 5.62 |
| **Cluster 2 (2577 voxels)** |  |  |  |  |  |
| Right intraparietal sulcus (IPS) | 36 | -54 | 46 |  | 7.52 |
| Right IPS | 38 | -46 | 46 | hIP3 | 7.42 |
| Right IPC | 60 | -44 | 20 | PFm | 7.25 |
| Right superior temporal gyrus (STG) | 36 | -54 | 46 |  | 5.6 |
| Right IPS | 30 | -62 | 52 | hIP3 | 5.21 |
| Right IPS | 32 | -68 | 32 | hIP5 | 5.19 |
| Right IPC | 46 | -34 | 42 | PFt | 5.08 |
| Right superior temporal sulcus (STS) | 50 | -28 | -2 |  | 5.02 |
| Right STS | 52 | -24 | -4 |  | 5.0 |
| Right IPS | 32 | -66 | 40 | hIP6 | 4.82 |
| **Cluster 3 (2292 voxels)** |  |  |  |  |  |
| Left IPC | -60 | -48 | 30 | PFcm | 7.74 |
| Left IPS | -40 | -48 | 48 | hIP3 | 7.73 |
| Left IPS | -46 | -40 | 44 |  | 7.51 |
| Left IPS | -24 | -66 | 50 | hIP6 | 7.51 |
| Left IPC | -58 | -44 | 36 | PF | 7.32 |
| Left IPS | -26 | -66 | 36 | hIP8 | 7.19 |
| Left IPS | -30 | -54 | 46 | hIP3 | 6.25 |
| Left angular gyrus | -56 | -52 | 12 |  | 5.36 |
| **Cluster 4 (1924 voxels)** |  |  |  |  |  |
| pMFC | 2 | 16 | 48 |  | 8.41 |
| pMFC | 6 | 26 | 36 |  | 8.35 |
| Superior frontal gyrus (SFG) | 10 | 14 | 60 | Area 6d2 | 6.13 |
| Pre-supplementary motor area (preSMA) | -4 | 4 | 56 | Area 6mr / preSMA | 5.75 |
| preSMA | 4 | 2 | 64 | Area 6mr / preSMA | 4.4 |
| SFG | 12 | 6 | 66 | Area 6d2 | 4.24 |
| **Cluster 5 (1050 voxels)** |  |  |  |  |  |
| Left IFJ | -44 | 6 | 32 |  | 8.36 |
| Left DLPFC | -40 | 28 | 24 |  | 4.55 |
| Left DLPFC | -40 | 30 | 24 |  | 4.55 |
| Left DLPFC | -38 | 36 | 36 |  | 3.72 |
| **Cluster 6 (1005 voxels)** |  |  |  |  |  |
| Left aI | -32 | 20 | 2 | Area Id7 | 8.42 |
| **Cluster 7 (538 voxels)** |  |  |  |  |  |
| Left lateral occipital cortex | -44 | -68 | -6 |  | 5.94 |
| Left fusiform gyrus | -42 | -78 | -8 | Area FG2 | 3.79 |
| **Cluster 8 (535 voxels)** |  |  |  |  |  |
| Right caudate nucleus | 12 | 8 | 6 |  | 7.72 |
| Right mediodorsal thalamus | 10 | -8 | 6 |  | 6.43 |
| Right caudate nucleus | 14 | 0 | 12 |  | 5.89 |
| Right putamen | 18 | 8 | 4 |  | 4.19 |
| Right pallidum | 14 | 4 | 0 |  | 3.58 |
| **Cluster 9 (334 voxels)** |  |  |  |  |  |
| Left dorsal premotor cortex | -26 | -4 | 56 | Area 6d3 | 6.95 |
| **Cluster 10 (190 voxels)** |  |  |  |  |  |
| Left caudate nucleus | -12 | 6 | 8 |  | 5.66 |
| Left mediodorsal thalamus | -12 | -12 | 4 |  | 4.84 |
| **Cluster 11 (133 voxels)** |  |  |  |  |  |
| Right precuneus | 10 | -68 | 50 |  | 5.23 |

### 3.3 Commonalities and differences in convergence of brain activity between failed and successful interference resolution

The conjunction analysis across experiments reporting brain activity related to committing errors or succeeding in interference control, respectively, revealed significant commonalities in recruiting bilateral aI, aMCC/preSMA, and bilateral TPJ (Figure 3A). Stronger convergence for brain activity linked to commission errors was found in bilateral middle insula, left SFG, aMCC, PCC, left posterior thalamus, and right V1 (Fig. 3B, depicted in green). Conversely, stronger convergence for brain activity linked to successful interference control was observed in bilateral IFJ extending into DLPFC, bilateral aI, bilateral dPMC, bilateral IPS/SPL, left lateral occipital cortex (including MT+), left middle and right superior temporal gyrus, right superior frontal gyrus and, subcortically, right caudate and right anterior thalamus (Fig. 3B, depicted in blue).

**Figure 3:**
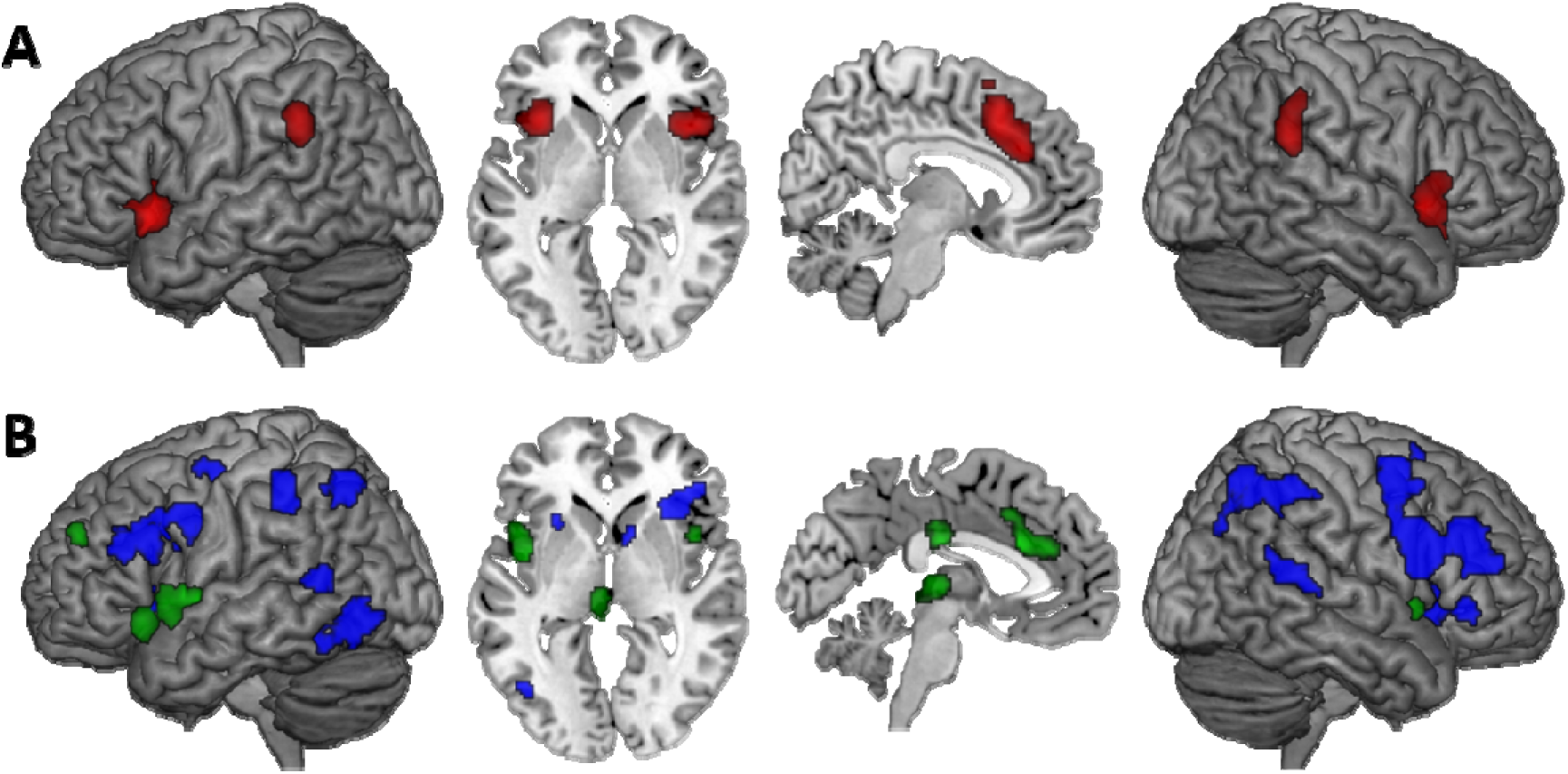
Commonalities and differences in erroneous versus successful responding were identified using (A) conjunction analysis across the results from the meta-analysis on commission errors and successful interference control respectively and (B) using meta-analytic contrast analysis between the two conditions.

In order to better visualise the conjoint effect for successful and failed interference resolution as well as the error-specific effect within the pMFC, Figure 4 shows the effects of the conjunction analysis (in red) as well as the specific effects for commission errors (in green) and successful interference control (in blue) in one overlay. The sections reveal that more posterior parts of pMFC are conjointly recruited during both failed and successful interference resolution, while the anterior portion features selectively stronger convergence for brain activity related to commission errors.

**Figure 4:**
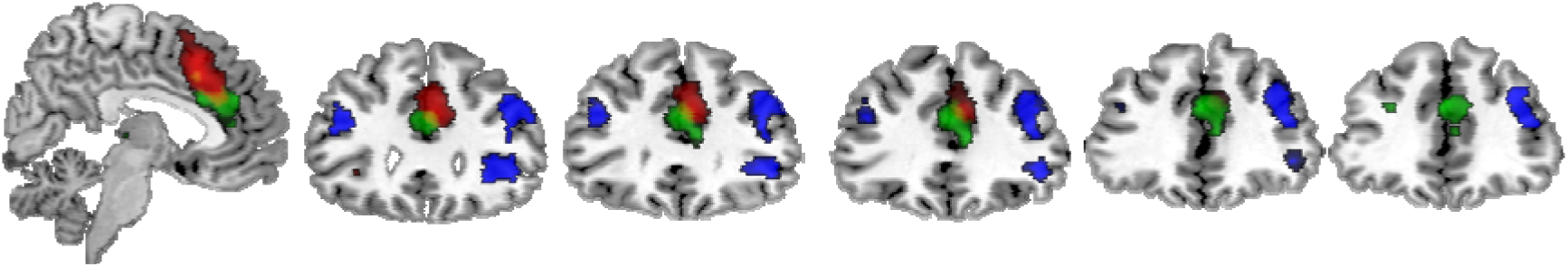
Overlay of the effects within the pMFC revealed by the conjunction analysis (in red) and the specific effects for commission errors (in green) and successful interference control (in blue) resulting from the meta-analytic contrast analysis.

Finally, main effects as well as conjunction and contrast analyses conducted with the reduced sample (i.e., only including publications that reported contrasts for both failed and successful interference control) are reported in detail in the supplementary material (Figure S1). Overall, the results were highly similar.

### 3.4 Control analyses separating between contrasting condition and type of task

#### 3.4.1 Meta-analyses on commission errors separated by contrasting condition

For the meta-analysis across experiments contrasting commission errors with low-level baseline or no-interference trials significant convergent activity within bilateral aI, a cluster located in the anterior aMCC, the preSMA, the dorsal PCC, and bilateral TPJ was found (Figure 5A). The meta-analysis across experiments contrasting commission errors with successful interference control trials revealed significant convergence within bilateral anterior and left middle insula, the aMCC/preSMA, right TPJ and, subcortically, the thalamus (Figure 5B). These additional analyses showed that the network consistently involved in error processing in general, i.e. when commission errors are compared to a no-interference or low-level condition, also revealed consistent increased activity when compared to a high-level condition, i.e. compared to successful interference trials.

**Figure 5:**
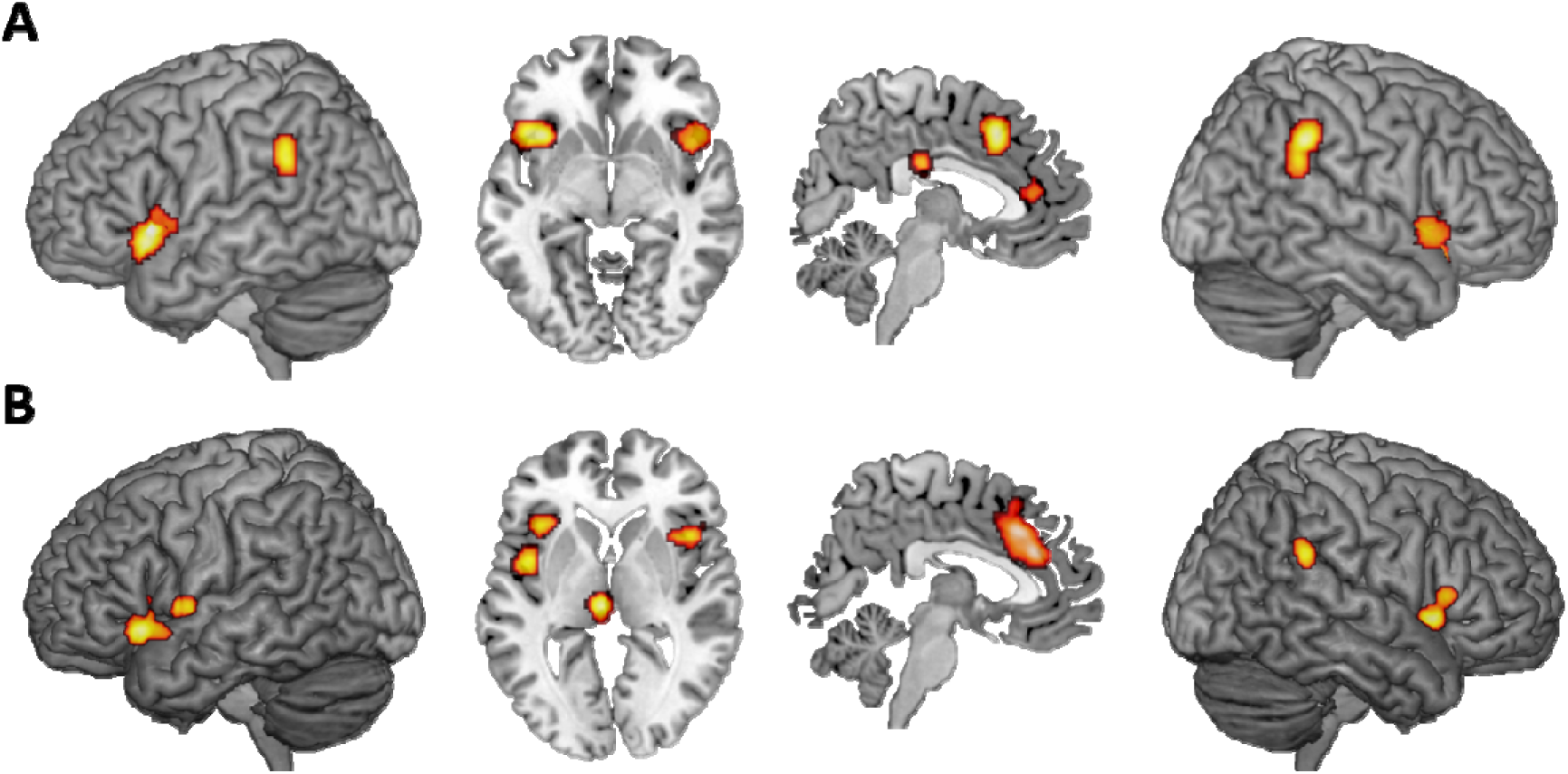
Control analyses investigating the effect of the contrasting condition by performing separate meta-analyses on experiments contrasting commission errors with no-interference or low-level baseline condition (A) and on experiments contrasting commission errors against successful interference control (B). All results are cluster-level p < .05 family-wise error-corrected for multiple comparisons, with a cluster forming threshold of p < .001 at the voxel level.

#### 3.4.2 Meta-analyses on error processing separated by type of task

To investigate if error processing in the classic response interference tasks and the classic response inhibition tasks may differ from each other, we additionally performed two meta-analyses focusing on task type. For the meta-analyses separated by task type, we not only included experiments contrasting commission errors versus successful interference control/no-interference/low-level baseline, but also experiments that did not model specifically commission errors but included commission as well as omission errors in the regressor of interest. This was done as there were not enough experiments focusing specifically on commission errors in the Stroop, Flanker, Simon or antisaccade tasks to calculate a robust meta-analysis. However, as in interference tasks most errors occur in the incongruent condition, but in the congruent condition omission errors are rarely observed, this should not have an impact on the results. As can be seen in Figure 6 both meta-analyses revealed a very similar set of regions, including bilateral aI, pMFC, TPJ and, subcortically the posterior thalamus.

**Figure 6:**
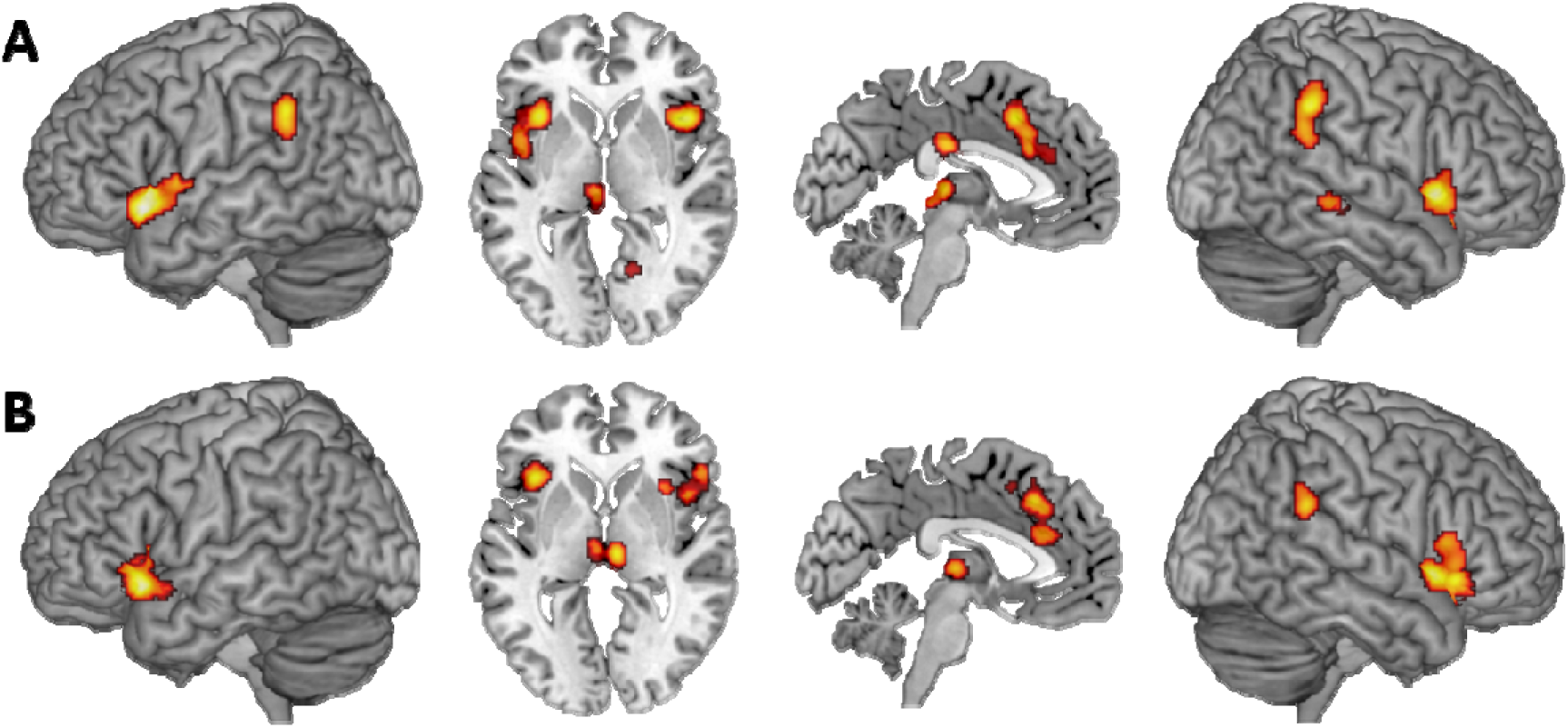
Control analyses investigating the effect of the type of paradigm by performing separate meta-analyses on error processing in the classic response inhibition tasks, i.e. go/no-go and stop signal tasks (A) and on error processing in the classic response interference tasks, i.e. Stroop, flanker, Simon, antisaccade tasks (B). All results are cluster-level p < .05 family-wise error-corrected for multiple comparisons, with a cluster forming threshold of p < .001 at the voxel level.

## 4. Discussion

We used coordinate-based ALE meta-analyses to examine the neural correlates of committing errors in experimental tasks that require the suppression of a predominant but inappropriate action and, in some cases, the concurrent initiation and execution of a context-appropriate alternative. Such commission errors are usually considered failures in top-down cognitive control, and characterising their neural underpinnings, we aimed to gain further insights into the neural architecture of intentional action control.

### 4.1 Common neural underpinnings of failed and successful interference control

Conjunction analyses were performed to isolate those brain regions that are consistently involved in performance monitoring irrespective of successful or erroneous responding. Commission errors as well as successful interference control revealed conjoint convergence in pMFC, bilateral aI and bilateral TPJ.

#### 4.1.1 Posterior medial frontal cortex

One of the most intensely discussed findings in the neuroscientific literature on error processing is the involvement of the pMFC. The discussion started with the seminal finding of Niki and Watanabe (1979) that individual monkey cingulate neurons show increased firing during commission errors. Later this was corroborated in human ERP studies, that identified the pMFC as the neural generator of the ERN (e.g. Debener et al., 2005; Miltner et al., 2003; Ullsperger and von Cramon, 2001). Combined EEG and fMRI recordings revealed that a greater ERN correlates with increased pMFC activity on the single-subject level (Debener et al., 2005; Iannocone et al., 2015). However, further research soon indicated that not only errors drive activation in this region but potentially some characteristics that errors have in common with other stimuli and conditions. For example, just the context of high error probability activates the pMFC, irrespective of actual error commission (Brown and Braver, 2005; Magno et al., 2006). Furthermore, neutral cues signalling a need for change in strategy also lead to pMFC activation, regardless of correct or erroneous responding (Amiez et al., 2005). Likewise, increased pMFC activity can be found in high-conflict trials when top-down control is low, such as incongruent trials that are preceded by congruent ones (e.g., Carter et al., 2000; Botvinick et al., 1999). The same behaviour was recently shown by single-cell recordings in rodents that performed a stop-change task (Bryden et al., 2019). pMFC firing was highest in stop trials that were preceded by go trials and decreased on trials following stop trials when rodents adapted their behaviour. Different models trying to explain the functional significance of pMFC activity have been proposed and tested formally (for recent reviews on pMFC models, see, e.g., Brockett and Roesch, 2021; or Vassena et al., 2017). While the various hypotheses differ in what exactly is signalled by the pMFC, there is general agreement that pMFC engagement contributes to performance monitoring. Specifically, it is believed to modulate attention in downstream regions and adaptation of control processes so that actions can be adjusted when the context requires it or errors occur (e.g. (MacDonald et al, 2000; Ullsperger and von Cramon, 2001; Ridderinkhof et al., 2004; Brockett et al., 2020; for a review, see Brockett and Roesch, 2021)). In line with this, Danielmeier and colleagues (Danielmeier et al., 2011; 2015) showed that pMFC activity during error trials is predictive of activity enhancement in sensory areas coding task-relevant information as well as activity suppression in task-irrelevant visual areas. Further, error-related pMFC activity is correlated with post-error adaptation processes, in particular PES (e.g. Garavan et al., 2002; Kerns et al., 2004). Supporting evidence from rodents reveals that pharmacological inactivation of pMFC eliminates PES (Narayanan et al., 2013).

In the present study, the conjunction analysis revealed common involvement of a cluster covering aMCC and preSMA for correct as well as erroneous responding, but the meta-analytic contrast analysis further revealed specific convergence of the anterior portion of aMCC for commission errors. The general pattern of a large aMCC/preSMA cluster being commonly involved in interference resolution irrespective of the accuracy of the behavioural outcome and the more anterior aMCC being specifically involved in erroneous responding is in line with previous fMRI studies that tested this difference directly (e.g. Ullsperger and van Cramon, 2001; Garavan et al., 2003; Kiehl et al., 2000). As cingulate subregions themselves and adjacent parts of the dorsomedial frontal cortex show distinct cytoarchitectonic profiles and connectivity (cf. e.g. Vogt et al., 2005; Neubert et al., 2015; Jin et al., 2018), the extended aMCC/preSMA cluster likely does not mediate one single process. Interestingly, recent single neuron recordings in patients revealed that not only in the aMCC but also the preSMA there are different neurons specifically signaling error or conflict, respectively (Fu et al., 2019). It thus appears that conflict detection and error monitoring are separable on the level of single neurons, but both processes can be localized within the aMCC and preSMA. This goes along with EEG evidence pointing towards a common underlying neuronal source within the pMFC for the ERN as well as the frontocentral N2 that can be observed prior to correct conflict trial responses (e.g. van Veen and Carter, 2002; Gruendler et al., 2011). In addition, specific involvement of the most anterior portion of the aMCC in error processing agrees with previous evidence pointing towards a specific role of anterior aMCC in negative affect related to performance slips, distuingshable from control functions related to conflict processing that can be localized more posteriorly (cf. Nee et al., 2011).

In conclusion, we argue that different subregions within the pMFC are associated with related but different functions in the context of performance monitoring. While the specific involvement of anterior aMCC in erroneous responding most likely reflects affective responses after error commission, more posterior parts of pMFC including aMCC and preSMA may mediate adaptive cognitive control processes necessary for successfully responding to high-conflict trials or after committing an error (Bush et al., 2000; Mohanty et al., 2007; van Veen and Carter, 2002).

#### 4.1.2 Insular cortex

The anterior part of the insular cortex revealed conjoint involvement in successful as well as erroneous responding. Moreover, while the insular cluster from the conjunction and contrast analyses showed some overlap, there was also a functional differentiation within the insula. In particular, the superior part of the aI in the right hemisphere showed stronger convergence for successful interference resolution, while inferior aI as well as more posterior parts of the insular cortex revealed stronger convergence for commission errors. The aI together with the pMFC forms the so-called saliency network (Seeley et al., 2007), which mediates switching between the DMN and executive control network (Sridharan et al., 2008). In particular, the aI is believed to respond to behaviourally salient stimuli or events and, through switching between the DMN and executive control network, facilitates necessary additional processing and control adjustments (Menon and Uddin, 2010). In the context of error processing, fMRI studies revealed stronger aI activity for aware compared to unaware errors (Hester et al., 2009, Klein et al., 2007) arguing for a crucial role of the insula in mediating awareness that an error has happened (for a review, see Ullsperger et al., 2010). Evidence from meta-analytic (Kurth et al., 2010) as well as resting-state fMRI data (Cauda et al., 2011; Kelly et al., 2012; Deen et al., 2011) provides evidence for a functional gradient within the insular cortex, with superior aI being involved in attentional and cognitive processes, inferior aI mediating emotional and autonomic processes, and posterior parts being involved in somatosensory processing. We hence conclude that the insula may support different functions during performance monitoring. Consistent involvement of the aI in both successful and erroneous responding may be explained by an increased salience of both interference and error trials, signaling the need to invest effort. While in trials with successful performance, the identification of the task-relevant stimulus property and the subsequent activation of cognitive control processes to guide behaviour occurs in time, in error trials, the detection of the error itself most likely leads to a recruitment of cognitive control strategies that facilitate correct performance in the following trial.

More inferior parts of the aI that revealed stronger convergence in error trials may reflect an autonomic response to errors. It has long been shown that the autonomic nervous system is sensitive to errors, as indicated by a decelerated heart rate after committing an error (Danev and de Winter, 1971; Fiehler et al., 2004; van der Veen et al., 2004), enlarged pupil dilation (Critchley et al., 2005; Wessel et al., 2011), or skin conductance responses (O’Connell et al., 2007). The latter study showed that only aware, but not unaware errors elicited a strong skin conductance response. Likewise, Wessel and colleagues (2011) found pupil dilation and post-error heart rate deceleration to be present in consciously perceived errors only. As the inferior part of the insula is associated with interoceptive awareness (Critchley et al., 2004), this region may mediate autonomic responses to errors, possibly contributing to error awareness (for reviews, see Ullsperger et al., 2010 and Klein et al., 2013).

The more posterior part of the insula, on the other hand, has been linked to the perception of somatosensory and proprioceptive signals, and evidence from patient studies points towards a role for motor and somatosensory awareness (Karnath et al., 2005; Spinazzola et al., 2008). Stronger convergence for error trials in posterior insular parts may hence be explained by the integration of motor feedback information of making the wrong button press.

Taken together, we argue that the insula mediates different functions in the context of performance monitoring and error processing. The superior portion of aI may have an active role in recruiting additional cognitive resources whenever needed to meet task demands, be it conflict trials or erroneous responses. In contrast, inferior aI and more posterior parts of the insula most likely contribute to the awareness that an error had occured. While inferior aI may integrate autonomic responses to errors, more posterior parts would integrate motor and somatosensory information about giving an erroneous behavioural response.

#### 4.1.3 Temporoparietal junction

A third region we found to be consistently involved in correct responding as well as commission errors is bilateral TPJ. This region is part of the ventral attention network and is associated with reorienting attention towards behaviourally relevant stimuli and events (Corbetta and Shulman, 2002; Corbetta and Shulman, 2008). These processes might also help in recovering from momentary lapses in attention through recruitment of attentional resources. In line with that, Weissman and colleagues (2006) found that greater target-evoked activity in the right TPJ was associated with better performance (i.e. faster RTs) in the next trial. Geng and Vossel (2013) argued for a more general role of the TPJ in contextual updating by evaluating and integrating stimulus information with internal models of task performance and expectations. Both attention reorientation as well as contextual updating play a key role in integrating stimulus information with task goals and adjustments of top-down expectations in conflict trials when the automatically activated predominant response tendency needs to be suppressed with concurrent initiation of the context-appropriate response. While these mechanisms might occur too late in error trials, they may support successful performance in the next trial. Interestingly, studies focusing on error awareness found increased activity for aware compared to unaware errors within the TPJ (e.g. Hester et al., 2005; 2012). This effect can be well integrated with the view that the TPJ mediates contextual updating as this process would especially come into play for consciously perceived errors compared to those that go unnoticed.

Importantly, consistent involvement of bilateral TPJ was found in the present study even though previously a right-lateralization of TPJ activity was observed for top-down action control (Langner et al., 2018), in line with the right TPJ’s hypothesised role in attentional reorienting (for a review, see Corbetta and Shulman, 2002). However, bilateral TPJ responses to task-relevant stimuli have been found in other studies as well (e.g. Downar et al., 2000; Kincade et al., 2005; for a discussion on bilateral TPJ involvement, see Geng and Vossel, 2013; Vossel et al., 2014). Likewise, resting-state as well as meta-analytic connectivity modelling studies have shown strong interhemispheric connectivity between left and right TPJ (Bzdok et al., 2013), questioning the theory of a strong right-lateralization. We conclude that bilateral TPJ most likely supports correct task performance through contextual updating during conflict trials when stimulus information needs to be integrated with the current task goal to respond in a non-dominant fashion. While in error trials these processes may occur too late, they may support correct behaviour in the following trial.

### 4.2 Regions specifically involved in error processing

Meta-analytic contrast analyses revealed a more strongly consistent involvement of PCC, left SFG, posterior thalamus, and V1 in erroneous than correct responding during cognitive interference tasks.

#### 4.2.1 Posterior cingulate cortex

Functionally, the PCC is considered part of the default-mode network, together with the ventromedial PFC and inferior parietal cortex, and typically shows deactivation during the performance of externally directed tasks but increased activity during the so-called “resting state” (Raichle et al., 2001; Buckner et al., 2008). As the DMN has typically shown activity anticorrelated to the dorsal fronto-parietal network (Fox et al., 2005), it has often been labelled the “task-negative” network. This characterization, however, ignores the DMN’s potentially active role in cognition (cf. Pearson et al., 2011). In particular, increased DMN activity has been observed in situations requiring internally directed attention, such as daydreaming, planning for the future, or autobiographical memory retrieval (Spreng and Grady, 2010).

Anatomical evidence suggests that the PCC is not a homogenous region (Vogt et al., 2009). Consistent error-related activity in our meta-analysis was found only within the dorsal part of PCC. Resting-state functional connectivity analyses in humans as well as monkeys show differential coupling patterns for dorsal versus ventral PCC (Leech et al., 2011; 2012; Margulies et al., 2009). While the ventral part is functionally fully integrated in the DMN, the dorsal PCC shows positive FC with a range of networks, including DMN but also the fronto-parietal control network, parts of the dorsal attention network, the sensorimotor network, and the salience network (Leech et al., 2012). Hence, the dorsal PCC may play a crucial role in linking functionally distinct networks that are recruited in a coordinated fashion when cognitive control is needed (cf. Leech and Sharp, 2014). In line with this notion, there is evidence for a more active role of the PCC when cognitive control demands increase. In particular, lesions of the PCC result in deficits in the implementation of new strategies as well as plan-following strategies in multitasking (cf. Burgess et al., 2000). Single-cell recordings in monkeys during a learning task showed phasic responses of PCC neurons to errors, with their activity being modulated by small rewards and novel stimuli. Moreover, reversible inactivation of PCC neurons impaired learning specifically in the most challenging condition, when monkeys were presented novel, low-value scenes, whereas PCC inactivation did not affect performance on well-learned associations (Heilbronner and Platt, 2013). Hence, PCC might be actively involved in detecting the need for enhanced cognitive engagement or goal-directed top-down modulation (Pearson et al., 2009), particularly coming into play when performance is poor. This, in turn, suggests that increased dorsal PCC firing is a consequence of poor performance, rather than a cause (cf. Heilbronner and Platt, 2013). This interpretation is in line with a recent fMRI study using a categorization learning experiment that required participants to identify the correct stimulus characteristics for categorization based on post-response feedback (Wolff and Brechmann, 2022). Here, negative (vs. positive) feedback elicited stronger activity in a widespread neural network including dorsal PCC. Importantly, dorsal PCC was the only region where this effect was enhanced in the revearsal learning phase when response contingencies switched. Dorsal PCC therefore most likely responds to feedback information signaling the need for subsequent behavioral adaptations. The dorsal PCC may then interact with the aMCC, which in turn implements corrective behavior (cf. Agam et al., 2011; Charles et al., 2013).

Taken together, even though PCC activity has rarely been discussed in neurobiological models of error processing, we argue that besides pMFC and aI, the dorsal PCC plays a key role in the coordinated recruitment of higher-order cognitive control. Convergence of dorsal PCC activity in the present study, therefore, most likely points to an active role of this region in post-error processing, by evaluating internal feedback from erroneous responses and passing on signals (potentially via the aMCC) that carry the need for behavioural adaptation.

#### 4.2.2 Superior frontal gyrus

The left SFG (MNI coordinate -28, 50, 28) was the only region in lateral PFC that showed consistent error-related activity. Interestingly, this cluster was located more superior and anterior to the clusters consistently showing involvement in successful interference resolution (cf. Fig. 3B). A host of studies have implicated lateral PFC in cognitive adaptation processes taking place after committing an error and preparing for high-conflict trials. However, there is considerable variability with regard to the exact neuroanatomical location and left versus right hemispheric involvement (e.g., MacDonald et al., 2000; Kerns et al., 2004). Evidence from the Stroop paradigm points towards a specific role of the left DLPFC in contexts requiring a temporal up-regulation of the attentional set (for a review, see Vanderhasselt et al., 2009). Crucially, however, these attentional adjustments were based on participants’ expectancies regarding the nature of the upcoming trial, but not on the amount of previous conflict per se. In contrast, the right DLPFC was hypothesised to support micro-adjustments in a moment-to-moment manner in conflict-driven contexts, leading to an up-regulation of cognitive control processes. Egner and Hirsch (2005), differentiating effects of conflict detection and adaptation, found a region in the left SFG, very close to our cluster, to be associated with the reduction in current-trial conflict due to exerting increased cognitive control after high-conflict trials. Along the same lines, Magno and colleagues (Magno et al., 2006) used a paradigm that allowed participants to reject a trial to avoid an error and found the same region in the left SFG to be more strongly activated for reject versus error trials and also a correlation of this region’s activity with performance. They hence concluded that this region is central to maintaining a representation of relevant task goals and the implementation of necessary changes in behaviour to avoid errors. Moreover, a recent rTMS study showed that inhibiting the left, but not right, DLPFC results in a reduced Pe component in a subsequently performed Go/No-Go task. As the ERN component was not affected, the study provides evidence for a left-sided DLPFC involvement in later stages of error-related processing (Masina et al., 2019). In conclusion, we would argue that the left SFG cluster we observed in our error-specific analysis reflects this region’s recruitment when top-down control is to be strengthened or focused to re-adapt cognitive processing to (again) meet behavioural goals after an error has been made.

#### 4.2.3 Posterior thalamus

Performance monitoring relies on the integrated functioning of the medial frontal cortex with the lateral PFC and the basal ganglia (Ullsperger et al., 2014b). Previous evidence suggests that within the thalamus, especially the ventral anterior (VA) and ventrolateral anterior (VLa) parts are strongly anatomically connected with the aMCC, and lesions affecting the VA/VLa nuclei lead to an abolished ERN (Seifert et al., 2011). Interestingly, the present meta-analysis revealed a consistent involvement of bilateral posterior thalamus extending into the mediodorsal thalamus in erroneous responding, and the meta-analytic contrast analysis provided further evidence for an error specifity of the posterior thalamus encompassing also the pulvinar nucleus. The pulvinar receives inputs from visual cortices (Arcaro et al., 2015; Shipp et al., 2003), and human lesion studies revealed a crucial role of the pulvinar in attentional selection (Danziger et al., 2004). As the pulvinar is also strongly connected to the TPJ in humans (Rosenberg et al., 2009), convergent activity during erroneous responding may reflect the interplay between the TPJ and pulvinar for post-error adjustments of attentional reorienting to the behaviourally relevant stimulus information.

Further, the error-specific cluster in the posterior thalamus may also include the lateral habenula (LHb), which plays an important role in signalling negative rewards or feedback (Shepard et al., 2006; Salas et al., 2010; Ullsperger and von Cramon, 2003; Baker et al., 2016). Single-cell recording in non-human primates as well as Granger causality and mediation analyses in humans argue for a regulatory role of the LHb on outcome-related signals in ventral tegmental area during error processing (Matsumoto and Hikosaka, 2007; Ide and Li, 2011). In particular, it is believed that negative feedback, such as an error, is accompanied by increased LHb activity, which in turn inhibits dopaminergic midbrain nuclei and hence decreases dopamine output (cf. Shepard et al., 2006). Kawai and colleagues (2015) recorded neural activity of LHb and pMFC in monkeys during a reversal learning task and found LHb to signal current negative outcomes, whereas pMFC neurons stored information from several past trials and signalled behavioural adjustment in the next trial.

While experiments with higher spatial resolution will be necessary to further disentangle the specific roles of thalamic nuclei for error processing and adaptation processes, we argue that across different interference paradigms particularly the posterior thalamus is recruited when errors occur. The habenular complex may take an active role in signalling ongoing negative outcomes and the pulvinar participating in post-error adjustment processes that may help to reorient attention to task-relevant information.

#### 4.2.4 V1

One reason for committing errors is the automation allocation of selective attention to distracting stimulus features, while task-relevant information is neglected or processed too late (Danielmeier et al., 2011). Evience from fMRI as well as MEG studies showed for instance that correct responding in the stop signal task depends on the strength of processing of the task-relevant (stop) stimulus (Boehler et al., 2009; 2010). Hence, after committing an error, a reallocation of attention to task-relevant information is required to enable correct performance in the next trial. fMRI studies have provided evidence for this top-down reallocation of attention after commission errors. For example, a study using the face-gender classification version of the Simon task found enhanced target processing in face-sensitive sensory cortex after commission errors (King et al., 2010). Likewise, activity increases in task-relevant visual areas have been observed in post-error trials in interference tasks, while activity in visual areas representing task-irrelevant distracting information decreased, and these modulations were correlated with error-related pMFC activity (Danielmeier et al., 2011; 2015). While the neural circuits underlying this top-down attentional modulations are not completely understood, a recent study using a 5-choice serial reaction time task in mice revealed a direct projection from the anterior cingulate area to the visual cortex selectively engaged in the post-error regulation of selective attention (Norman et al., 2021).

While the present study provides evidence for a specific and consistent involvement of right V1 during incorrect responding, the functional significance of this effect has rarely been discussed in the respective papers. Interestingly, a recent study investigating momentary lapses of attention in a multisensory environment found not only increased pre-stimulus activity in the DMN, but also that increased pre-stimulus activity within a region of the right calcarine cortex close to our cluster led to impaired subsequent task performance as indicated by increased RT (Su et al., 2020). We hence argue that the stronger convergence of right V1 activity during error trials may reflect an automatic, bottom-up driven allocation of attention to irrelevant stimulus properties (in the context of interference tasks) or a stronger processing of the go stimulus (in the context of inhibition tasks), interfering with the processing of task-relevant information and resulting in committing errors.

Besides V1, we did not find convergence in any other region that one might have expected to be associated with error-prone states, such as an increased activity in the DMN. While increased activity in the DMN has been found in trials preceding errors (Weissmann et al., 2006; Su et al., 2020), due to the low temporal resolution of fMRI we would have expected to also find consistent activity in these regions when an error occurs. One reason for the lack of convergence in regions related to error-prone states may be that errors can result from different causes, such as attentional lapses or motor control failures (van Driel et al., 2012; Weissman et al., 2006; Su et al. 2020) as well as maladaptation of cognitive control (Steinhauser et al., 2012) or insufficient task rule activation (Possin et al., 2009), hence reducing the probability to find convergence across studies. Further, there may be task-specific error-prone states, as errors in the stop-signal task, where participants are usually pushed towards a 50% accuracy rate, may be driven by different factors than errors in a flanker task, where errors are much more infrequent and more likely result from short-time attentional lapses and control failures. However, our meta-analyses separating between paradigm classes, i.e. the classic response interference tasks and the classic response inhibition tasks, did show a very similar set of regions consistently recruited during erroneous responding (cf. Fig. 6A,B). As these meta-analyses included only a relatively small number of studies, additional data is needed to more thoroughly investigate task-specific, or more generally, error-type-specific effects.

### 4.3 Regions specifically involved in successful conflict resolution

The broad fronto-parietal network consistently involved in correct interference resolution (Fig. 2) well replicates results of previous studies. However, it does not seem to be involved selectively in interference control processes but rather also in other facets of executive functioning. As it responds in a very domain-general manner across a diversity of tasks probing executive functioning, it has been labelled the multiple-demand network (Duncan and Owen, 2000, Duncan et al., 2010; Müller et al., 2015) or cognitive control network (Cole and Schneider, 2007). As discussed above, parts of the network (i.e., pMFC, bilateral aI and TPJ) were also involved in error-related processing as revealed by the conjunction analysis. In contrast, posterior parietal, superior and lateral prefrontal parts of the network were more strongly involved in correct interference control. The lack of consistent activation within these frontoparietal regions during error trials may reflect the failure to activate or apply the correct stimulus–response contingencies for successful response execution and thus suggest a mechanism for how errors come about. This notion is also supported by studies showing that the mere anticipation of interference (such as a cue indicating a high probability of a stop signal) is enough to activate this network (Jahfari et al., 2012; Smittenaar et al., 2013), suggesting its importance for the correctness of behavioural output.

While there may be many roads to errors, here we highlight two possible sources based on the networks involved in successful conflict resolution. The dorsal parietal and superior frontal regions of the successful conflict resolution network are part of a dorsal attention network that has been consistently involved in visuospatial attention and goal-directed selection of stimulus-response associations (e.g. Corbetta and Shulman 2002; Corbetta et al., 2008; Cieslik et al., 2010). This network controls voluntary attention by sending top-down control signals to lower-level sensory areas, thereby facilitating the processing of task-relevant stimuli and suppression of irrelevant information (Corbetta et al., 2000; Hopfinger et al., 2000; Vossel et al., 2014). The dorsal attention network increases activity when the number or priority of distractors presented in the vicinity of the target stimulus is increased (Lanssens et al., 2020; Molenberghs et al., 2008, Nobre et al, 2003; Wei et al., 2019). Likewise, stimulation of the superior frontal cortex as well as the IPS modulates the ability of filtering out distractors (Lega et al., 2019; Moos et al., 2012; Jigo et al., 2018). One road that may therefore lead to errors are attention control failures, as distracting stimuli can impair stopping on the stop-signal task (Verbruggen, Stevens, & Chambers, 2014), likely impacting appropriate cognitive control. Consistently less activity in error trials within the dorsal attention network may therefore reflect failures in suppressing task-irrelevant information (such as flanking distractors in the flanker task) and/or delayed processing of the task-relevant information (such as the stop-signal is processed too late), leading to the execution of the automatic, overlearned behaviour before the correct alternative could be initiated.

Next, the lPFC has been assumed to subserve the planning of behavioural responses and encoding of task relevant rule sets (Tanji and Hoshi, 2001; Mian et al., 2014). Similarly, non-human primate studies provide evidence for the lPFC playing a key role in the representation of task rules that link relevant stimulus features and appropriate motor responses (Hoshi and Tanji, 2004; Ninokura et al., 2004; White and Wise, 199, Sakamoto et al., 2022). In humans, right lPFC atrophy is associated with increased rule violation errors (Possin et al., 2009). The inferior frontal junction (IFJ), a region contained within the cluster found in lPFC, has been proposed to continuously reactivate relevant task rules that link relevant stimulus features and non-dominant responses (Cieslik et al. 2015). The lack of convergence in the lPFC for incorrect compared to correct responding may thus indicate momentary lapses in task rule activation resulting in the lack of suppression of the predominant response and its subsequent execution. As we did not find convergence within any other brain regions that we would hypothesize to be associated with error-prone states apart from V1, errors may not simply result from an overactivation of task-irrelevant regions but more likely from an insufficient activation of the task-relevant regions. Future studies may test this hypothesis by using the current results as seed regions to investigate their activation level prior to an erroneous response and when it happens. This may help to obtain a better understanding of the temporal BOLD fluctuations within the network and test the hypothesis that commission errors go along with reduced activity in the dorsal attention network and lPFC, as a sign of attentional control failures or short-time lapses in task-rule reactivation.

### 4.4 Limitations

Importantly, already over a decade ago, Grinband and colleagues (2008; 2011) argued that pMFC activity may not reflect conflict or error likelihood at all but rather is sensitive to trial-to-trial differences in response times. As incongruent trials are commonly associated with longer RTs than congruent ones, this may be a confound in fMRI studies not adjusting for differences in RTs and may also impact the observed activation level in other task-positive brain regions (Yarkoni et al., 2009; see also recent preprints investigating this effect: Beldzik and Ullsperger, 2023 (preprint); Mumford et al., 2023 (preprint)). As most studies investigating interference resolution do not adjust for RT differences, this may also have an impact on the present meta-analytic findings. However, as commission errors are usually associated with shorter RTs than high-conflict trials (e.g. Ridderinkhof et al., 2003), the regions showing common involvement across erroneous and successful responding are most likely not strongly affected by this effect. However, future studies investigating interference control should specifically take into account the effect of condition differences in response speed and how these might modulate observed activation levels in the task network.

### 4.5 Conclusion

This study provides evidence that monitoring one’s behavioural performance relies on the integrated functioning of different neural networks. The pMFC and aI, central hubs of the salience network (Seeley et al., 2007), are consistently involved during correct as well as erroneous responding and most likely the key players in the evaluation of situations calling for increased cognitive control, such as high-conflict trials or errors. Meta-analytic contrast analyses revealed some functional differentiation within these regions, which may point towards the specific salience of errors as failures may be particularly relevant for behavioural learning and adaptation.

Besides the well-established roles of pMFC and insula, a set of other regions was also found to be consistently involved in error-related processing. In particular, the dorsal PCC, which has been less commonly assigned a key role in the context of error monitoring, revealed specific convergence for erroneous responding. We therefore argue that this region works in concert with pMFC and aI, coming into play particularly when internal monitoring indicates that the performance level is (too) low, by subserving the evaluation of internal and external feedback from erroneous responses and forwarding the need for behavioural adaptation processes via the pMFC to the SFG, which then implements subsequent top-down behavioral control.

Bilateral TPJ, which was consistently activated across successful and erroneous responding, likely mediates contextual updating in high-conflict trials that require the integration of stimulus information with current task goals. While these processes may occur too late in error trials, they likely support correct task performance in subsequent ones, thereby contributing to post-error adaptation processes.

The dorsal attention network and lPFC, on the other hand, showed stronger convergence for successful response execution, which may point towards a failure to inhibit distracting sensory information and insufficient task-rule reactivation in erroneous trials. Together with the fact that besides V1 we did not find convergence in any other region whose increased activity could be assumed to reflect error-prone states, it seems that the processes leading to skill-based slips and lapses according to Reasonśs nomenclature (Reason, 1990) can most likely be localised within the network responsible for correct cognitive action control itself, rather than outside the network.

## Acknowledgements

We thank all contacted authors who contributed results of relevant contrasts not explicitly reported in the original publications, and we apologise to all authors whose eligible papers we might have missed.

This study was supported by the National Institute of Mental Health (R01-MH074457) and the Helmholtz Portfolio Theme “Supercomputing and Modeling for the Human Brain”.

This project has received funding from the European Research Council (ERC) under the European Union’s Horizon 2020 research and innovation programme (grant agreement No 101018805).

