## Supplementary data for "Success versus failure in cognitive control: meta-analytic evidence from neuroimaging studies on error processing"

### Figure S1: Flowchart providing an overview of meta-analytic workflow

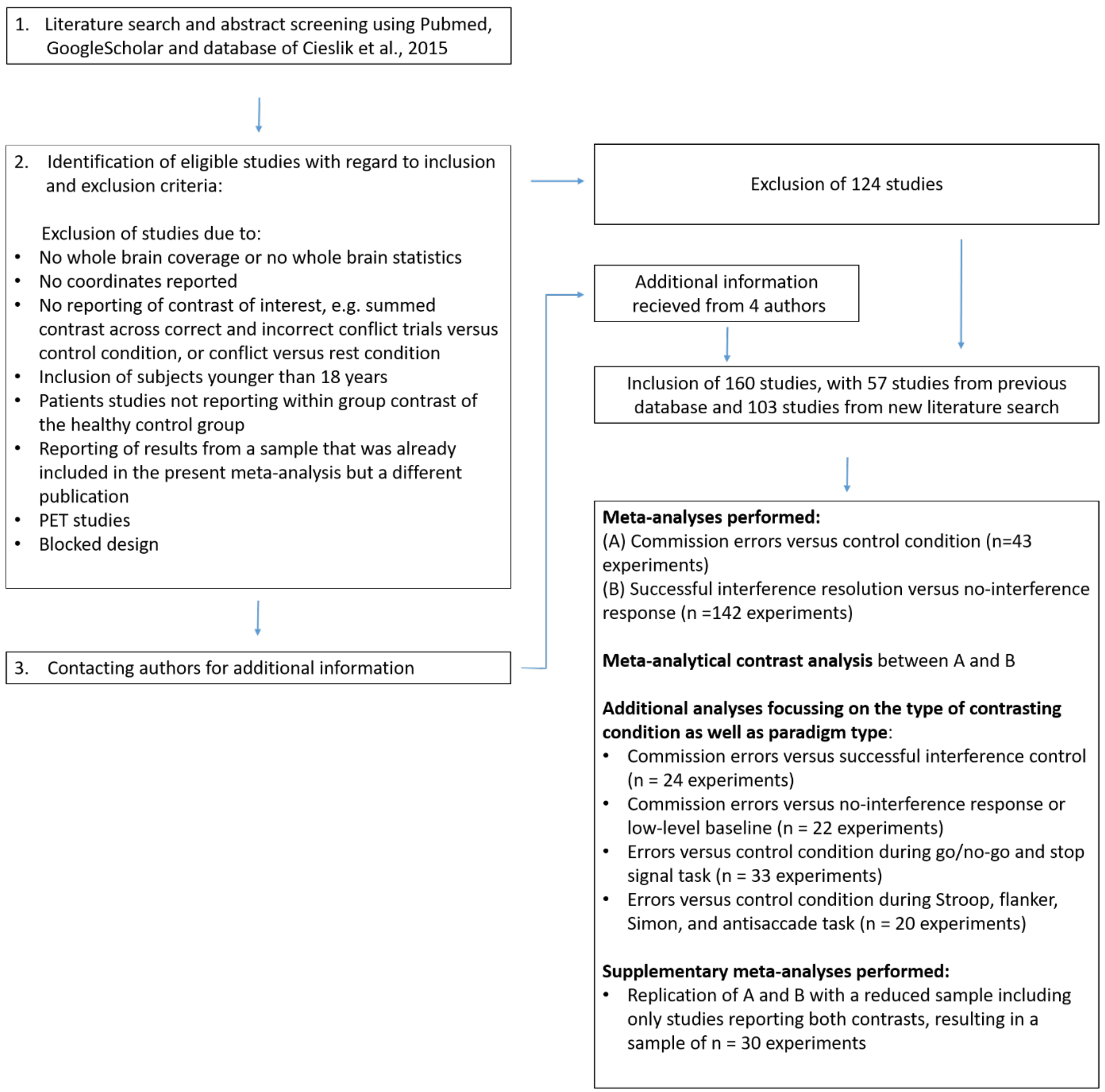

**Supplementary analyses:**

**Meta-analyses on the reduced sample, i.e. only including publications that reported contrasts for both, commission errors and successful interference control:**

The meta-analysis on the reduced sample of commission errors revealed significant convergent activity in bilateral aI, left SFG, the pMFC, PCC, bilateral TPJ, right superior temporal sulcus, and left posterior thalamus (Figure S1A). The meta-analysis on the reduced sample of successful interference control revealed significant convergence in bilateral aI, right IFJ and adjacent lPFC, left precentral gyrus, bilateral TPJ and IPS (Figure S1B).

The minimum conjunction analysis across commission errors and successful interference control revealed common involvement of bilateral aI, aMCC/preSMA as well as bilateral TPJ (Figure S3C). Stronger convergence for commission errors was found in the middle insula, left SFG, aMCC, dorsal PCC, and left posterior thalamus (Figure S1D in green). In contrast, stronger convergence for successful conflict resolution was revealed in right IFJ, right middle frontal gyrus, left precentral gyrus, and right IPS/IPC (Figure S1D in blue).

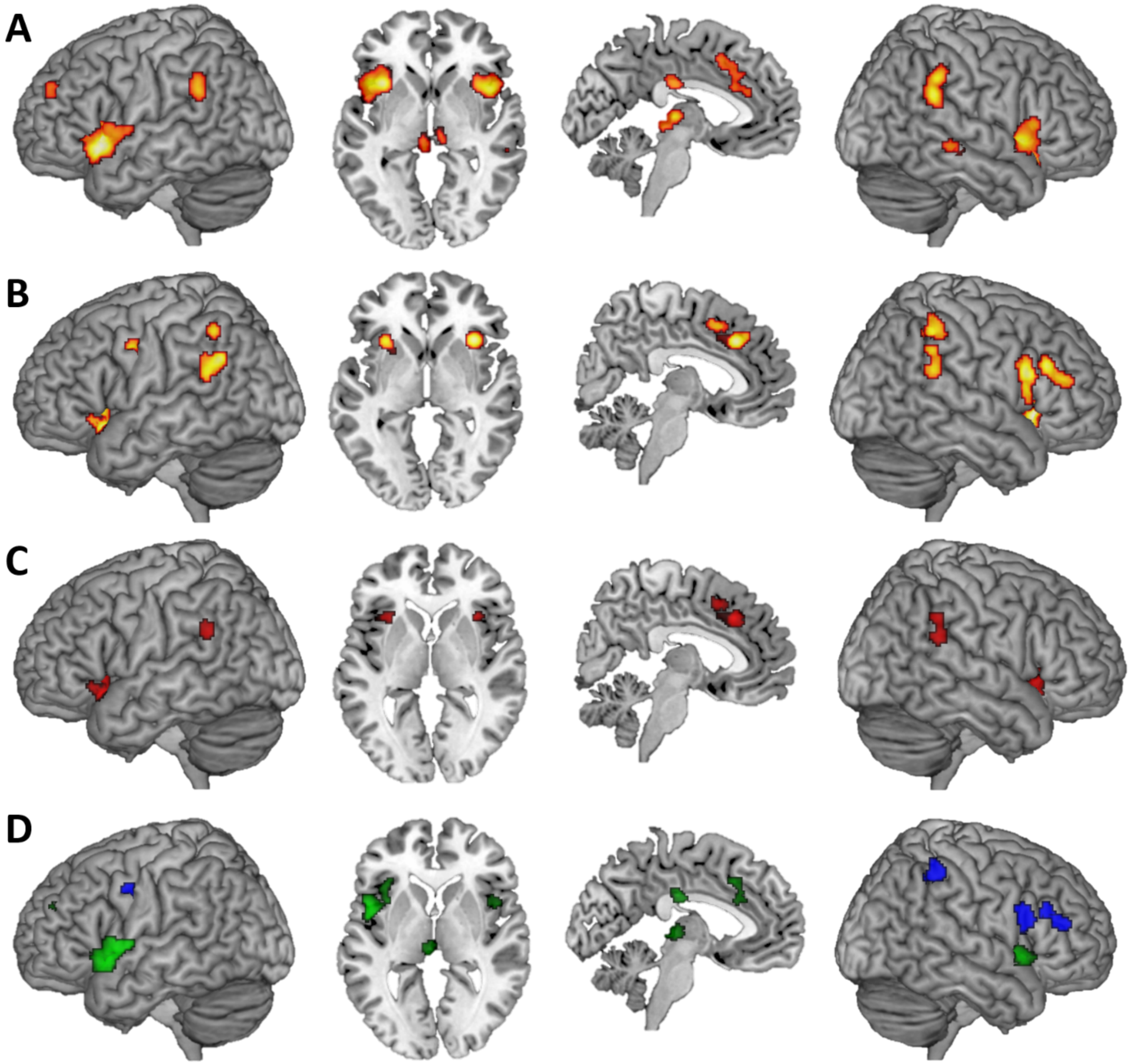
A
B
C
D

**Checklist for neuroimaging meta-analyses**

| | |
|---|---|
| The research question is specifically defined | .YES, commonalities and differences in erroneous versus successful interference control performing the following meta-analyses:<br>1) Commission errors<br>2) Successful interference control<br>3) Meta-analytic contrast:<br>Commission errors versus successful interference control<br><br>4) Control analyses separating between contrasting condition<br>A) Commission errors (versus low-level baseline or no-interference control)<br>B) Commission errors (versus successful interference control)<br><br>5) Control analyses separating between paradigm classes<br>A) Errors during classic response inhibition tasks<br>B) Errors during classic interference tasks<br><br>6) Supplementary contrast analysis with reduced, matched sample contrasting commission errors versus successful interference control |
| The literature search was systematic | YES, it included the following keywords in the following databases:<br><br>1) "Commission error", "error", "erroneous responding", "Stroop", flanker", "antisaccade" "stimulus response compatibility", "go/no-go", "stop signal", "Simon", "interference", "inhibition", "fMRI", "neuroimaging".<br>2) Databases: PubMed (https://www.ncbi.nlm.nih.gov/pubmed/) and Google Scholar (http://scholar.google.de) |
| Detailed inclusion and exclusion criteria are included | YES, and reasons of non-standard criterion were: |

| | |
|---|---|
| | Inclusion of:<br><br>- fMRI studies, which reported the coordinates in a standard reference space (Montreal Neurological Institute (MNI) or Talairach-Tournoux system (TAL))<br>- Healthy participants over the age of 18 and without any pharmacological manipulations<br>- Only activation data, no deactivation data included<br>- Whole-brain data<br>- Only event-related designs, blocked-designs excluded<br>- No correlation or interaction with other variables (e.g., performance measures) |
| Sample overlap was taken into account | YES, using the following method:<br><br>- If a study reported several experiments eligible for inclusion, the reported coordinates were pooled to constitute a single experiment<br>- If a study separately reported two or more subject groups, e.g. young and old participants with separate results, the coordinates were not pooled<br>- If different studies reported results of the same sample of participants only one of the study was included |
| All experiments use the same search coverage (state how brain coverage is assessed and how small volume corrections and conjunctions are taken into account) | YES, the search coverage is the following:<br><br>- Only whole-brain coverage<br>- Exclusion of ROI studies<br>- Exclusion of partial brain coverage |
| Studies are converted to a common reference space | YES, using the following conversion(s):<br><br>- Coordinates reported in Talairach space were converted to MNI space (Lancaster et al., 2007). |

| | |
|---|---|
| Data extraction have been conducted by two investigators (ideal case) or double checked by the same investigator (state how double-checking was performed) | YES, the following authors:<br><br>- Edna C. Cieslik, checked inclusion criteria, extracted coordinate information as well as other information regarding number of participants, space, task type, specific contrast, control condition<br>- Edna Cieslik double-checked data from Cieslik et al., 2015 for inclusion in the present study due to more stringent inclusion criteria in the present meta-analysis (e.g. only inclusion of event-related design and correct successful interference control)<br>- All extracted information was double-checked by research assistants |
| The paper includes a table with at least the references, basic study description (e.g. for fMRI tasks, stimuli), contrasts and basic sample descriptions (e.g. size, mean age and gender distribution, specific characteristics) of the included studies, source of information (e.g. contact with authors), reference space | YES, the table includes the following information:<br><br>- First author and year<br>- Number of subjects<br>- Space<br>- Control Condition<br>- Source of coordinates<br>- If further information was received by the authors<br>- How coordinates were treated (MNI or Talairach) when space was not clearly specified in original study |
| The study protocol and all analyses was planned beforehand, including the methods and parameters used for inference, correction for multiple testing, etc. | YES:<br><br>1) No non-planned or post-hoc analyses.<br>2) The meta-analysis used the default methods and parameters of our group |
| The meta-analysis includes diagnostics | Contributions are provided in the supplementary material |

**Contributions:**

**Meta-analysis on commission errors:**

**40 experiments contributing to Cluster 1 (Posterior medial frontal cortex):**

| | | | | |
|---|---|---|---|---|
| Agam et al., 2011 | Chevrier et al., 2007 | Debener et al., 2005 | Fassbender et al., 2004 | Fiehler et al., 2004 |
| Garavan et al., 2002 | Garavan et al., 2003 | Ham et al., 2013 | Hester et al., 2004b | Hughes et al., 2012 |
| Hughes et al., 2013 | Iannaccone et al., 2015 | Kiehl et al., 2000 | Klein et al., 2007 | Li et al., 2008 |
| Lütcke and Frahm, 2007 | Matthews et al., 2005 | Menon et al., 2001 | Hester et al., 2005 | Ramautar et al., 2006 |
| Rubia et al., 2003 | Sharp et al., 2010 | Sosic-Vasic et al., 2012 | Steele et al., 2014 | |
| Ullsperger and von Cramon, 2001 | | Wittfoth et al., 2008 | Dambacher et al., 2015 | Hough et al., 2016 |
| Ko et al. 2014 | Köhler et al., 2018 | Boecker et al., 2011 | Jahfari, 2012 | Mohammadi et al., 2015 |
| Xu et al., 2017 | Chen et al., 2015 | Jahfari, 2011 | Li et al., 2006 | Krönke et al., 2018 |
| Holmes et al., 2009 | Boehler et al., 2010 | | | |

**33 experiments contributing to Cluster 2 (left anterior insula):**

| | | | | |
|---|---|---|---|---|
| Agam et al., 2011 | Debener et al., 2005 | Fassbender et al., 2004 | Garavan et al., 2002 | Ham et al., 2013 |
| Hester et al., 2004b | Hughes et al., 2012 | Iannaccone et al., 2015 | Klein et al., 2007 | Li et al., 2008 |
| Lütcke and Frahm, 2007 | Menon et al., 2001 | Hester et al., 2005 | Ramautar et al., 2006 | Sosic-Vasic et al., 2012 |
| Steele et al., 2014 | Ullsperger and von Cramon, 2001 | | Wittfoth et al., 2008 | Dambacher et al., 2015 |
| Hough et al., 2016 | Hsu et al., 2017 | Ko et al. 2014 | Köhler et al., 2018 | Boecker et al., 2011 |
| Jahfari, 2012 | Mohammadi et al., 2015 | Xu et al., 2017 | Chen et al., 2015 | Jahfari, 2011 |
| Li et al., 2006 | Krönke et al., 2018 | Holmes et al., 2009 | Boehler et al., 2010 | |

**27 experiments contributing to Cluster 3 (right anterior insula):**

| | | | | |
|---|---|---|---|---|
| Fassbender et al., 2004 | Garavan et al., 2002 | Hughes et al., 2012 | Iannaccone et al., 2015 | Klein et al., 2007 |
| Li et al., 2008 | Menon et al., 2001 | Ramautar et al., 2006 | Sosic-Vasic et al., 2012 | Steele et al., 2014 |

Ullsperger and von Cramon, 2001  
Wittfoth et al., 2008  
Dambacher et al., 2015  
Hough et al., 2016  
Hsu et al., 2017  
Ko et al. 2014  
Köhler et al., 2018  
Boecker et al., 2011  
Harle, 2016  
Jahfari, 2012  
Mohammadi et al., 2015  
Xu et al., 2017  
Chen et al., 2015  
Jahfari, 2011  
Krönke et al., 2018  
Holmes et al., 2009  
Boehler et al., 2010

**21 experiments contributing to Cluster 4 (right inferior parietal cortex/temporoparietal junction):**

Chevrier et al., 2007  
Fassbender et al., 2004  
Garavan et al., 2002  
Ham et al., 2013  
Hester et al., 2004b  
Hughes et al., 2012  
Klein et al., 2007  
Hester et al., 2005  
Rubia et al., 2003  
Sosic-Vasic et al., 2012  
Steele et al., 2014  
Ullsperger and von Cramon, 2001  
Dambacher et al., 2015  
Hough et al., 2016  
Köhler et al., 2018  
Boecker et al., 2011  
Chen et al., 2015  
Jahfari, 2011  
Krönke et al., 2018  
Holmes et al., 2009  
Boehler et al., 2010

**16 experiments contributing to Cluster 5 (posterior thalamus):**

Garavan et al., 2002  
Hester et al., 2004b  
Li et al., 2008  
Lütcke and Frahm, 2007  
Ramautar et al., 2006  
Sosic-Vasic et al., 2012  
Steele et al., 2014  
Wittfoth et al., 2008  
Dambacher et al., 2015  
Köhler et al., 2018  
Jahfari, 2012  
Li et al., 2006  
Krönke et al., 2018  
Holmes et al., 2009  
Boehler et al., 2010  
Zandbelt et al., 2011

**14 experiments contributing to Cluster 6 (left inferior parietal cortex/temporoparietal junction):**

Chevrier et al., 2007  
Fassbender et al., 2004  
Ham et al., 2013  
Hester et al., 2004b  
Hughes et al., 2012  
Klein et al., 2007  
Ramautar et al., 2006  
Sosic-Vasic et al., 2012  
Steele et al., 2014  
Ullsperger and von Cramon, 2001  
Dambacher et al., 2015  
Boecker et al., 2011  
Jahfari, 2012  
Jahfari, 2011

**9 experiments contributing to Cluster 7 (left superior frontal gyrus):**

| | | | | |
|---|---|---|---|---|
| Debener et al., 2005 | Ham et al., 2013 | Iannaccone et al., 2015 | Kiehl et al., 2000 | Steele et al., 2014 |
| Dambacher et al., 2015 | Jahfari, 2012 | Chen et al., 2015 | Boehler et al., 2010 | |

**12 experiments contributing to Cluster 8 (posterior cingulate cortex):**

| | | | | |
|---|---|---|---|---|
| Chevrier et al., 2007 | Fassbender et al., 2004 | Hester et al., 2004b | Klein et al., 2007 | Matthews et al., 2005 |
| Steele et al., 2014 | Wittfoth et al., 2008 | Boecker et al., 2011 | Jahfari, 2012 | Xu et al., 2017 |
| Jahfari, 2011 | Boehler et al., 2010 | | | |

**8 experiments contributing to Cluster 9 (primary visual cortex):**

| | | | | |
|---|---|---|---|---|
| Agam et al., 2011 | Li et al., 2008 | Steele et al., 2014 | Dambacher et al., 2015 | Köhler et al., 2018 |
| Li et al., 2006 | Holmes et al., 2009 | Boehler et al., 2010 | | |

**Meta-analysis on successful interference resolution:**

**118 experiments contributing experiments to Cluster 1 (right anterior insula / inferior frontal junction / dorsolateral prefrontal cortex):**

| | | | | |
|---|---|---|---|---|
| Garavan et al., 1999 | Liddle et al. , 2001 | | | |
| Ullsperger et von Cramon, 2001 | | Garavan et al., 2002 | | |
| Garavan et al., 2003 | Ye et al., 2009 | Cieslik et al., 2010 | Wittfoth et al., 2008 | Chikazoe et al., 2007 |
| Aarts et al., 2008 | Lee et al., 2008 | King et al., 2012 | Frühholz et al., 2011 | Basten et al., 2011 |
| Becker et al., 2008 | Christensen et al., 2011_1 | Grandjean et al., 2012 | Kim et al., 2011 | Kim et al., 2012 |
| Kerns et al., 2005 | Luks et al., 2007 | Rubia et al., 2006 | van Eijk et al., 2015 | van Eijk et al., 2015_2 |
| Prakash et al., 2009 B | Chen et al., 2018 A | Chen et al., 2018 B | Hough et al., 2016 | Huang et al., 2012 |
| Lesh et al., 2013 | Luethi et al., 2016 | Manard et al., 2017 | Piai et al., 2013 | Pompei et al., 2011 |
| Verstynen et al., 2014 | Jiang et al., 2014 | Jaspar et al., 2014 | Overbeek et al., 2019 | |
| Purmann & Pollmann, 2015 | Shin & Kim, 2015 | Ramm et al., 2020 | Kim et al., 2014 | Krönke et al., 2018 |
| Schmidt et al., 2012 | Wallentin et al., 2015 | Cojan et al., 2015 | Bunge et al., 2002 | Dichter et al., 2007 |
| Holtmann et al., 2013 | Korsch et al., 2014 | Korsch et al., 2014 | Paschke et al., 2015 | Trautwein et al., 2016 |
| Berron et al., 2015 | Chuang et al., 2014 | Daamen et al., 2015 | Ivanov et al., 2012 | Chevrier et al., 2007 |
| Kelly et al., 2004 | Chikazoe et al., 2009b | Marco-Pallarés et al., 2008 | Nakata et al., 2008 | Zheng et al., 2008 |
| Cai and Leung, 2009 | Boehler et al., 2010 | Sharp et al., 2010 | Walther et al., 2010 | Rothmayr et al., 2011 |
| Aron and Poldrack, 2006 | Cai and Leung, 2011 | Chikazoe et al., 2009a | Hughes et al., 2012 | Jahfari et al., 2011 |
| Kenner et al., 2010 | Tabu et al., 2012 | Xue et al., 2008 | Hester et al., 2004a | Hester et al., 2004b |
| Kaladjian et al., 2009a | Fauth-Bühler et al., 2012 | Tabu et al., 2011 | Berkmann et al., 2014 | Boecker et al., 2011 |
| Cai et al. 2014 | Coxon et al., 2016 | Ganos et al., 2014 | Ghahremani et al., 2012 | Harle et al., 2016 |
| Hughes et al., 2013 | Jahfari et al., 2012 | Jahfari et al., 2015 | Ko et al., 2016 | Lavallee et al., 2014 |
| Lenartowicz et al. 2010 | Lorenz et al., 2015 | Mohammadi et al., 2015 | Rae et al., 2014 | |
| Rodriguez-Pujadas et al., 2014 | Schel et al., 2014 | Sebastian et al., 2013a | Sebastian et al., 2013b | Sebastian et al., 2013b_2 |
| Sebastian et al., 2017 | Swann et al. 2012 | Van der Meer et al., 2013 | Wilbertz et al., 2014 | Xu et al., 2015 |
| Xu et al., 2017 | Zandbelt and Fink, 2010 | Zandbelt et al. 2011 | Chen et al., 2015 | Fedota et al., 2015 |
| Fuentes-Claramonte et al., 2016 | | Hsu et al., 2017 | Köhler et al., 2018 | O'Connor et al., 2012 |
| Steele et al. 2014 | Sebastian et al., 2012 | | | |

**106 experiments contributing to Cluster 2 (right intraparietal sulcus):**

| | | | | |
|---|---|---|---|---|
| Garavan et al., 1999 | Liddle et al. , 2001 | Ullsperger et von Cramon, 2001 | | Garavan et al., 2002 |
| Garavan et al., 2003 | Cieslik et al., 2010 | Wittfoth et al., 2008 | Chikazoe et al., 2007 | Aarts et al., 2008 |
| King et al., 2012 | Basten et al., 2011 | Grandjean et al., 2012 | Kim et al., 2011 | Kim et al., 2012 |
| Page et al., 2009 | Kerns et al., 2005 | Luks et al., 2007 | Chen et al., 2018 B | Hough et al., 2016 |
| Huang et al., 2012 | Manard et al., 2017 | Pompei et al., 2011 | Verstynen et al., 2014 | Jiang et al., 2014 |
| Coderre et al., 2013 | Overbeek et al., 2019 | Shin & Kim, 2015 | Kim et al., 2014 | Krönke et al., 2018 |
| Schmidt et al., 2012 | Wallentin et al., 2015 | Cojan et al., 2015 | Bunge et al., 2002 | Dichter et al., 2007 |
| Holtmann et al., 2013 | Iannaccone et al., 2015 | Paschke et al., 2015 | Trautwein et al., 2016 | Chuang et al., 2014 |
| Daamen et al., 2015 | Ivanov et al., 2012 | Kelly et al., 2004 | Chikazoe et al., 2009b | |
| Marco-Pallarés et al., 2008 | Nakata et al., 2008 | Zheng et al., 2008 | Cai and Leung, 2009 | Boehler et al., 2010 |
| Sharp et al., 2010 | Walther et al., 2010 | Rothmayr et al., 2011 | Aron and Poldrack, 2006 | Chikazoe et al., 2009a |
| Fassbender et al., 2004 | Hughes et al., 2012 | Jahfari et al., 2011 | Kenner et al., 2010 | Tabu et al., 2012 |
| Xue et al., 2008 | Hester et al., 2004a | Hester et al., 2004b | Kaladjian et al., 2007 | Kaladjian et al., 2009a |
| Mazzola-Pomietto et al., 2009 | Kaladjian et al., 2009b | Fauth-Bühler et al., 2012 | Berkmann et al., 2014 | Boecker et al., 2011 |
| Cai et al. 2014 | Coxon et al., 2016 | Coxon et al., 2016_2 | Ganos et al., 2014 | Ghahremani et al., 2012 |
| Harle et al., 2016 | Hughes et al., 2013 | Jahfari et al., 2012 | Jahfari et al., 2015 | Ko et al., 2016 |
| Lavallee et al., 2014 | Lenartowicz et al. 2010 | Lorenz et al., 2015 | Mohammadi et al., 2015 | Montojo et al., 2013 |
| Rae et al., 2014 | Rodriguez-Pujadas et al., 2014 | Schel et al., 2014 | Sebastian et al., 2013a | Sebastian et al., 2013b |
| Sebastian et al., 2013b_2 | Sebastian et al., 2017 | Swann et al. 2012 | Van der Meer et al., 2013 | Wilbertz et al., 2014 |
| Xu et al., 2015 | Xu et al., 2017 | Zandbelt and Fink, 2010 | Zandbelt et al. 2011 | Czapla et al., 2017 |
| Fedota et al., 2015 | Fuentes-Claramonte et al., 2016 | Köhler et al., 2018 | Meffert et al., 2016 | O'Connor et al., 2012 |
| Steele et al. 2014 | Ramautar et al., 2006 | Sebastian et al., 2012 | | |

**103 experiments contributing to Cluster 3 (left inferior parietal cortex):**

| | | | | |
|---|---|---|---|---|
| Garavan et al., 1999 | Liddle et al. , 2001 | Garavan et al., 2003 | Ye et al., 2009 | Cieslik et al., 2010 |
| Chikazoe et al., 2007 | Aarts et al., 2008 | Hart et al., 2010 | King et al., 2012 | Frühholz et al., 2011 |
| Basten et al., 2011 | Becker et al., 2008 | Grandjean et al., 2012 | Kim et al., 2011 | Kim et al., 2012 |
| Page et al., 2009 | Kerns et al., 2005 | Luks et al., 2007 | Zurawska et al., 2011 | van Eijk et al., 2015 |
| Prakash et al., 2009 A | Prakash et al., 2009 B | Chen et al., 2018 A | Chen et al., 2018 B | Hough et al., 2016 |
| Huang et al., 2012 | Lesh et al., 2013 | Luethi et al., 2016 | Manard et al., 2017 | Pompei et al., 2011 |
| Verstynen et al., 2014 | Jiang et al., 2014 | Coderre et al., 2013 | Jaspar et al., 2014 | Overbeek et al., 2019 |

| | | | | |
|---|---|---|---|---|
| Purmann & Pollmann, 2015 | Shin & Kim, 2015 | Kim et al., 2014 | Krönke et al., 2018 | Schmidt et al., 2012 |
| Wallentin et al., 2015 | Cojan et al., 2015 | Bunge et al., 2002 | Dichter et al., 2007 | Holtmann et al., 2013 |
| Korsch et al., 2014 | Paschke et al., 2015 | Trautwein et al., 2016 | Chuang et al., 2014 | Daamen et al., 2015 |
| Ivanov et al., 2012 | Wang et al., 2014 | Kelly et al., 2004 | Chikazoe et al., 2009b | |
| Marco-Pallarés et al., 2008 | Zheng et al., 2008 | Cai and Leung, 2009 | Boehler et al., 2010 | Sharp et al., 2010 |
| Walther et al., 2010 | Rothmayr et al., 2011 | Chikazoe et al., 2009a | Hughes et al., 2012 | Jahfari et al., 2011 |
| Kenner et al., 2010 | Tabu et al., 2012 | Xue et al., 2008 | Hester et al., 2004a | Hester et al., 2004b |
| Kaladjian et al., 2007 | Kaladjian et al., 2009b | Fauth-Bühler et al., 2012 | Berkmann et al., 2014 | Boecker et al., 2011 |
| Cai et al. 2014 | Coxon et al., 2016 | Coxon et al., 2016_2 | Ganos et al., 2014 | Ghahremani et al., 2012 |
| Harle et al., 2016 | Jahfari et al., 2012 | Ko et al., 2016 | Lavallee et al., 2014 | Lorenz et al., 2015 |
| Rae et al., 2014 | Sebastian et al., 2013a | Sebastian et al., 2013b | Sebastian et al., 2013b_2 | Sebastian et al., 2017 |
| Swann et al. 2012 | Van der Meer et al., 2013 | Wilbertz et al., 2014 | Xu et al., 2015 | Xu et al., 2017 |
| Zandbelt and Fink, 2010 | Zandbelt et al. 2011 | Czapla et al., 2017 | Fuentes-Claramonte et al., 2016 | Köhler et al., 2018 |
| Meffert et al., 2016 | O'Connor et al., 2012 | Steele et al. 2014 | Sebastian et al., 2012 | |

**97 experiments contributing to Cluster 4 (posterior medial frontal cortex):**

| | | | | |
|---|---|---|---|---|
| Garavan et al., 1999 | Kiehl et al., 2000 | Liddle et al. , 2001 | Ullsperger et von Cramon, 2001 | |
| Garavan et al., 2002 | Ye et al., 2009 | Cieslik et al., 2010 | Chikazoe et al., 2007 | Aarts et al., 2008 |
| King et al., 2012 | Frühholz et al., 2011 | Basten et al., 2011 | Becker et al., 2008 | Grandjean et al., 2012 |
| Kim et al., 2011 | Kim et al., 2012 | Kerns et al., 2005 | Luks et al., 2007 | van Eijk et al., 2015 |
| van Eijk et al., 2015_2 | Chen et al., 2018 A | Chen et al., 2018 B | Huang et al., 2012 | Lesh et al., 2013 |
| Piai et al., 2013 | Verstynen et al., 2014 | Jiang et al., 2014 | Coderre et al., 2013 | Hinault et al., 2018 |
| Overbeek et al., 2019 | Purmann & Pollmann, 2015 | Shin & Kim, 2015 | Ramm et al., 2020 | Kim et al., 2014 |
| Krönke et al., 2018 | Schmidt et al., 2012 | Wallentin et al., 2015 | Cojan et al., 2015 | Bunge et al., 2002 |
| Dichter et al., 2007 | Holtmann et al., 2013 | Iannaccone et al., 2015 | Korsch et al., 2014 | Paschke et al., 2015 |
| Trautwein et al., 2016 | Berron et al., 2015 | Chuang et al., 2014 | Daamen et al., 2015 | Ivanov et al., 2012 |
| Kelly et al., 2004 | Chikazoe et al., 2009b | Marco-Pallarés et al., 2008 | Nakata et al., 2008 | Cai and Leung, 2009 |
| Boehler et al., 2010 | Sharp et al., 2010 | Walther et al., 2010 | Rothmayr et al., 2011 | Aron and Poldrack, 2006 |
| Chikazoe et al., 2009a | Jahfari et al., 2011 | Kenner et al., 2010 | Tabu et al., 2012 | Xue et al., 2008 |
| Hester et al., 2004a | Hester et al., 2004b | Tabu et al., 2011 | Berkmann et al., 2014 | Boecker et al., 2011 |

| | | | | |
|---|---|---|---|---|
| Cai et al. 2014 | Chikara et al., 2018 | Coxon et al., 2016 | Ganos et al., 2014 | Ghahremani et al., 2012 |
| Harle et al., 2016 | Jahfari et al., 2015 | Ko et al., 2016 | Lenartowicz et al. 2010 | Lorenz et al., 2015 |
| Mohammadi et al., 2015 | Rae et al., 2014 | Sebastian et al., 2013a | Sebastian et al., 2013b | Sebastian et al., 2013b_2 |
| Van der Meer et al., 2013 | Wilbertz et al., 2014 | Xu et al., 2015 | Xu et al., 2017 | Zandbelt and Fink, 2010 |
| Zandbelt et al. 2011 | Chen et al., 2015 | Fuentes-Claramonte et al., 2016 | Köhler et al., 2018 | O'Connor et al., 2012 |
| Steele et al. 2014 | Ramautar et al., 2006 | Sebastian et al., 2012 | | |

**70 experiments contributing to Cluster 5 (left inferior frontal junction / dorsolateral prefrontal cortex):**

| | | | | |
|---|---|---|---|---|
| Liddle et al. , 2001 | Ye et al., 2009 | Chikazoe et al., 2007 | King et al., 2012 | Basten et al., 2011 |
| Becker et al., 2008 | Grandjean et al., 2012 | Kim et al., 2011 | Kim et al., 2012 | Page et al., 2009 |
| Kerns et al., 2005 | Rubia et al., 2006 | van Eijk et al., 2015_2 | Prakash et al., 2009 A | Prakash et al., 2009 B |
| Chen et al., 2018 A | Huang et al., 2012 | Lesh et al., 2013 | Luethi et al., 2016 | Manard et al., 2017 |
| Pompei et al., 2011 | Verstynen et al., 2014 | Jiang et al., 2014 | Coderre et al., 2013 | Jaspar et al., 2014 |
| Overbeek et al., 2019 | Shin & Kim, 2015 | Ramm et al., 2020 | Kim et al., 2014 | Krönke et al., 2018 |
| Schmidt et al., 2012 | Wallentin et al., 2015 | Cojan et al., 2015 | Bunge et al., 2002 | Dichter et al., 2007 |
| Holtmann et al., 2013 | Paschke et al., 2015 | Trautwein et al., 2016 | Chuang et al., 2014 | Chikazoe et al., 2009b |
| Nakata et al., 2008 | Cai and Leung, 2009 | Boehler et al., 2010 | Chikazoe et al., 2009a | Fassbender et al., 2004 |
| Kenner et al., 2010 | Tabu et al., 2012 | Hester et al., 2004a | Hester et al., 2004b | Kaladjian et al., 2007 |
| Kaladjian et al., 2009a | Kaladjian et al., 2009b | Fauth-Bühler et al., 2012 | Berkmann et al., 2014 | Boecker et al., 2011 |
| Coxon et al., 2016 | Ko et al., 2016 | Lavallee et al., 2014 | Lorenz et al., 2015 | Mohammadi et al., 2015 |
| Rae et al., 2014 | Sebastian et al., 2013a | Sebastian et al., 2013b_2 | Wilbertz et al., 2014 | Xu et al., 2015 |
| Xu et al., 2017 | Zandbelt and Fink, 2010 | Zandbelt et al. 2011 | Köhler et al., 2018 | Meffert et al., 2016 |

**73 experiments contributing to Cluster 6 (left anterior insula):**

| | | | | |
|---|---|---|---|---|
| Ullsperger et von Cramon, 2001 | | Garavan et al., 2002 | Ye et al., 2009 | Cieslik et al., 2010 |
| Chikazoe et al., 2007 | Aarts et al., 2008 | Becker et al., 2008 | Christensen et al., 2011_1 | Grandjean et al., 2012 |
| Kim et al., 2012 | Kerns et al., 2005 | Luks et al., 2007 | Zurawska et al., 2011 | Rubia et al., 2006 |
| van Eijk et al., 2015_2 | Prakash et al., 2009 B | Chen et al., 2018 B | Hough et al., 2016 | Huang et al., 2012 |
| Manard et al., 2017 | Verstynen et al., 2014 | Jiang et al., 2014 | Jaspar et al., 2014 | |

| | | | | |
|---|---|---|---|---|
| Purmann & Pollmann, 2015 | Shin & Kim, 2015 | Kim et al., 2014 | Schmidt et al., 2012 | Wallentin et al., 2015 |
| Cojan et al., 2015 | Holmes et al., 2009 | Trautwein et al., 2016 | Kelly et al., 2004 | Chikazoe et al., 2009b |
| Cai and Leung, 2009 | Boehler et al., 2010 | Aron and Poldrack, 2006 | Chikazoe et al., 2009a | Fassbender et al., 2004 |
| Hughes et al., 2012 | Jahfari et al., 2011 | Kenner et al., 2010 | Tabu et al., 2012 | Hester et al., 2004a |
| Fauth-Bühler et al., 2012 | Tabu et al., 2011 | Berkmann et al., 2014 | Cai et al. 2014 | Congdon et al., 2014 |
| Coxon et al., 2016 | Ganos et al., 2014 | Ghahremani et al., 2012 | Jahfari et al., 2012 | Jahfari et al., 2015 |
| Ko et al., 2016 | Lorenz et al., 2015 | Mohammadi et al., 2015 | Rae et al., 2014 | Sebastian et al., 2013a |
| Sebastian et al., 2013b | Sebastian et al., 2013b_2 | Sebastian et al., 2017 | Van der Meer et al., 2013 | Wilbertz et al., 2014 |
| Xu et al., 2015 | Xu et al., 2017 | Zandbelt and Fink, 2010 | Zandbelt et al. 2011 | Chen et al., 2015 |
| Fuentes-Claramonte et al., 2016 | | Köhler et al., 2018 | O'Connor et al., 2012 | Steele et al. 2014 |
| Sebastian et al., 2012 | | | | |

**54 experiments contributing to Cluster 7 (left lateral occipital cortex):**

| | | | | |
|---|---|---|---|---|
| Garavan et al., 1999 | Kiehl et al., 2000 | Ullsperger et von Cramon, 2001 | | Ye et al., 2009 |
| Wittfoth et al., 2008 | Chikazoe et al., 2007 | Frühholz et al., 2011 | Basten et al., 2011 | Grandjean et al., 2012 |
| Rubia et al., 2006 | van Eijk et al., 2015 | van Eijk et al., 2015_2 | Chen et al., 2018 B | Hough et al., 2016 |
| Luethi et al., 2016 | Manard et al., 2017 | Jiang et al., 2014 | Coderre et al., 2013 | Shin & Kim, 2015 |
| Krönke et al., 2018 | Schmidt et al., 2012 | Wallentin et al., 2015 | Cojan et al., 2015 | Holtmann et al., 2013 |
| Iannaccone et al., 2015 | Korsch et al., 2014 | Trautwein et al., 2016 | Chuang et al., 2014 | Daamen et al., 2015 |
| Ivanov et al., 2012 | Kelly et al., 2004 | Chikazoe et al., 2009a | Marco-Pallarés et al., 2008 | Cai and Leung, 2009 |
| Boehler et al., 2010 | Cai and Leung, 2011 | Hester et al., 2004a | Kaladjian et al., 2007 | Kaladjian et al., 2009a |
| Kaladjian et al., 2009b | Fauth-Bühler et al., 2012 | Bobb et al., 2011 | Ko et al., 2016 | Schel et al., 2014 |
| Sebastian et al., 2013a | Sebastian et al., 2013b | Sebastian et al., 2013b_2 | Van der Meer et al., 2013 | Wilbertz et al., 2014 |
| Fedota et al., 2015 | Meffert et al., 2016 | O'Connor et al., 2012 | Steele et al. 2014 | Sebastian et al., 2012 |

**53 experiments contributing to Cluster 8 (right caudate nucleus and mediodorsal thalamus):**

| | | | | |
|---|---|---|---|---|
| Ullsperger et von Cramon, 2001 | | Garavan et al., 2002 | Chikazoe et al., 2007 | Aarts et al., 2008 |
| Hart et al., 2010 | Basten et al., 2011 | Christensen et al., 2011_1 | Grandjean et al., 2012 | Kim et al., 2011 |

| | | | | |
|---|---|---|---|---|
| Kim et al., 2012 | Rubia et al., 2006 | Chen et al., 2018 B | Hough et al., 2016 | Luethi et al., 2016 |
| Piai et al., 2013 | Verstynen et al., 2014 | Coderre et al., 2013 | Shin & Kim, 2015 | Krönke et al., 2018 |
| Schmidt et al., 2012 | Wallentin et al., 2015 | Korsch et al., 2014 | Berron et al., 2015 | Daamen et al., 2015 |
| Ivanov et al., 2012 | Chevrier et al., 2007 | Chikazoe et al., 2009b | Boehler et al., 2010 | Aron and Poldrack, 2006 |
| Cai and Leung, 2011 | Chikazoe et al., 2009a | Kenner et al., 2010 | Hester et al., 2004a | Hester et al., 2004b |
| Fauth-Bühler et al., 2012 | Berkmann et al., 2014 | Congdon et al., 2014 | Coxon et al., 2016 | Harle et al., 2016 |
| Lorenz et al., 2015 | Montojo et al., 2013 | Schel et al., 2014 | Sebastian et al., 2013a | Sebastian et al., 2013b |
| Sebastian et al., 2013b_2 | Wilbertz et al., 2014 | Xu et al., 2015 | Xu et al., 2017 | Zandbelt and Fink, 2010 |
| Zandbelt et al. 2011 | Fuentes-Claramonte et al., 2016 | | Hsu et al., 2017 | O'Connor et al., 2012 |

**36 experiments contributing to Cluster 9 (left dorsal premotor cortex):**

| | | | | |
|---|---|---|---|---|
| Liddle et al. , 2001 | Cieslik et al., 2010 | Chikazoe et al., 2007 | Aarts et al., 2008 | King et al., 2012 |
| Frühholz et al., 2011 | Grandjean et al., 2012 | Kerns et al., 2005 | Zurawska et al., 2011 | Jiang et al., 2014 |
| Coderre et al., 2013 | Jaspar et al., 2014 | Purmann & Pollmann, 2015 | Krönke et al., 2018 | Schmidt et al., 2012 |
| Wallentin et al., 2015 | Holtmann et al., 2013 | Iannaccone et al., 2015 | Trautwein et al., 2016 | Daamen et al., 2015 |
| Chikazoe et al., 2009a | Boehler et al., 2010 | Tabu et al., 2012 | Xue et al., 2008 | Hester et al., 2004a |
| Hester et al., 2004b | Fauth-Bühler et al., 2012 | Harle et al., 2016 | Lorenz et al., 2015 | Sebastian et al., 2013a |
| Sebastian et al., 2013b | Sebastian et al., 2013b_2 | Wilbertz et al., 2014 | Zandbelt and Fink, 2010 | Zandbelt et al. 2011 |
| Sebastian et al., 2012 | | | | |

**36 experiments contributing to Cluster 10 (right caudate nucleus and mediodorsal thalamus):**

| | | | | |
|---|---|---|---|---|
| Ullsperger et von Cramon, 2001 | | Garavan et al., 2002 | Chikazoe et al., 2007 | Aarts et al., 2008 |
| Basten et al., 2011 | Kim et al., 2011 | Kim et al., 2012 | Kerns et al., 2005 | Hough et al., 2016 |
| Huang et al., 2012 | Verstynen et al., 2014 | Coderre et al., 2013 | Shin & Kim, 2015 | Krönke et al., 2018 |
| Schmidt et al., 2012 | Wallentin et al., 2015 | Berron et al., 2015 | Daamen et al., 2015 | Chikazoe et al., 2009b |
| Boehler et al., 2010 | Aron and Poldrack, 2006 | Chikazoe et al., 2009a | Hester et al., 2004a | Hester et al., 2004b |
| Fauth-Bühler et al., 2012 | Berkmann et al., 2014 | Coxon et al., 2016 | Lorenz et al., 2015 | Schel et al., 2014 |
| Sebastian et al., 2013b_2 | Sebastian et al., 2017 | Xu et al., 2015 | Xu et al., 2017 | Zandbelt and Fink, 2010 |
| Fuentes-Claramonte et al., 2016 | | O'Connor et al., 2012 | | |

**27 experiments contributing to Cluster 11 (right precuneus):**

Chikazoe et al., 2007
Jiang et al., 2014
Daamen et al., 2015
Xue et al., 2008
Lorenz et al., 2015
Steele et al. 2014
Aarts et al., 2008
Coderre et al., 2013
Ivanov et al., 2012
Berkmann et al., 2014
Rae et al., 2014
Sebastian et al., 2012
Rubia et al., 2006
Wallentin et al., 2015
Chikazoe et al., 2009b
Bobb et al., 2011
Sebastian et al., 2013b_2
Huang et al., 2012
Holtmann et al., 2013
Marco-Pallarés et al., 2008
Coxon et al., 2016
Xu et al., 2017
Verstynen et al., 2014
Chuang et al., 2014
Chikazoe et al., 2009a
Coxon et al., 2016_2
Zandbelt and Fink, 2010

| Author | n | space | paradigm | correct/control | contrasted against | type of error | inhibition / interference | Source of coordinates |
|---|---|---|---|---|---|---|---|---|
| Agam et al., 2011 | 30 | TAL | antisaccade | correct | correct_inc | CE | class. interference | SM Table 3 |
| Barke et al., 2017 | 75 | MNI | flanker | control | correct_con_inc | GE | class. interference | Table 1 |
| Boecker et al., 2011 | 15 | MNI | stop signal task | control | general_go | CE | inhibition | SM Table 2 |
| Boehler et al., 2010 | 15 | MNI | stop signal task | control, correct | correct_go, correct_stop | CE | inhibition | SM Table 4, SM Table 3 |
| Chen et al., 2015 | 25 | MNI | go/no-go | control | general_go | CE | inhibition | Table 2 |
| Chevrier et al., 2007 | 14 | TAL | stop signal task | other | baseline | CE | inhibition | Table 1 |
| Critchley et al., 2005 | 15 | MNI | stroop | control | correct_con_inc_neu | GE | class. interference | Table 1 |
| Dambacher et al., 2015 | 15 | TAL | go/no-go | control | correct_go | CE | inhibition | Table 1 |
| Debener et al., 2005 | 13 | TAL | flanker | correct | correct_inc | CE | class. interference | Table 1 |
| Fassbender et al., 2004 | 18 | TAL | go/no-go | other | baseline_during_rest | CE | inhibition | Table 2 |
| Fiehler et al., 2004 | 27 | TAL | flanker | correct | correct_inc | CE | class. interference | Table 2 |
| Fitzgerald et al, 2005 | 7 | MNI | flanker | other | all_other | GE | class. interference | Table 1 |

| | | | | | | | | |
|---|---|---|---|---|---|---|---|---|
| Garavan et al., 2002 | 14 | TAL | go/no-go | control | baseline_go | CE | inhibition | Table 1 |
| Garavan et al., 2003 | 16 | TAL | go/no-go | control | baseline_go | CE | inhibition | Table 1 |
| Ham et al., 2013 | 35 | MNI | simon | control, correct | correct_con, correct_inc | GE | class. interference | Table 1, Table 1 |
| Harle, 2016 | 34 | TAL | stop signal task | control | correct_go | CE | inhibition | SM Table 3 |
| Hester et al., 2004b | 15 | TAL | go/no-go | control | baseline_go | CE | inhibition | Table 2 |
| Hester et al., 2005 | 13 | TAL | go/no-go | control | baseline_go | CE | inhibition | Table 1 |
| Holmes et al., 2010 | 33 | MNI | flanker | correct | correct_inc | CE | class. interference | Table 3 |
| Hough et al., 2016 | 22 | MNI | go/no-go | other | baseline | CE | inhibition | SM Table 5 |
| Hsu et al., 2017 | 20 | MNI | go/no-go | control | general_go | CE | inhibition | Table 3 |
| Hughes et al., 2012 | 10 | MNI | stop signal task | control | baseline_go | CE | inhibition | Table 4 |
| Hughes et al., 2013 | 15 | MNI | stop signal task | control | baseline_go | CE | inhibition | Table 3 |
| Iannaccone et al., 2015 | 15 | MNI | flanker | correct | correct_inc | CE | class. interference | Table 2 |
| Jahfari, 2011 | 20 | MNI | stop signal task | control | general_go | CE | inhibition | Table 4 |
| Jahfari, 2012 | 16 | MNI | stop signal task | control | general_go | CE | inhibition | Table 5 |
| Kerns et al., 2005 | 13 | TAL | stroop | control | correct_con | GE | class. interference | Table 1 |
| Kiehl et al., 2000 | 14 | MNI | go/no-go | control, correct | baseline_go, correct_stop | CE | inhibition | Table 1, Table 1 |
| King et al., 2010 | 21 | TAL | simon | control | correct_con_inc | GE | class. interference | Table 1 |
| Klein et al., 2007 | 13 | TAL | antisaccade | correct | correct_inc | CE | class. interference | SM Table A |
| Ko et al. 2014 | 23 | MNI | go/no-go | correct | correct_stop | CE | inhibition | Table 3 |
| Köhler et al., 2018 | 33 | MNI | go/no-go | correct | correct_stop | CE | inhibition | Table 1 |
| Krönke et al., 2018 | 118 | MNI | Stroop | correct | correct_inc | CE | class. interference | App. 2, Table 6 |

| | | | | | | | | |
|---|---|---|---|---|---|---|---|---|
| Li et al., 2006 | 24 | TAL | stop signal task | correct | correct_stop | CE | inhibition | Table 2 |
| Li et al., 2008 | 40 | MNI | stop signal task | correct | correct_stop | CE | inhibition | Table 1 |
| Lütcke and Frahm, 2008 | 11 | MNI | go/no-go | correct | correct_stop | CE | inhibition | Table 2 |
| Marco-Pallarés et al., 2008 | 10 | MNI | flanker | control | correct_con_inc | GE | class. interference | Table 1 |
| Matthews et al., 2005 | 16 | TAL | stop signal task | correct,<br>correct | correct_stop (short failed inhibit trials > short successful inhibit trials),<br>correct_stop (long failed inhibit trials > long successful inhibit trials) | CE<br>CE | inhibition | Table 2<br>Table 3 |
| Menon et al., 2001 | 12 | MNI | go/no-go | correct | correct_stop | CE | inhibition | Table 1 |
| Mohammadi et al., 2015 | 17 | MNI | stop signal task | control, correct | general_go, correct_stop | CE | inhibition | SM Table 3, SM Table 3 |
| Ramautar et al., 2006 | 16 | TAL | stop signal task | correct,<br>correct | correct_stop (unsuccessful stop > successful stop in low frequency stop signal condition),<br>correct_stop (unsuccessful stop > successful stop in high frequency stop signal condition) | CE<br>CE | inhibition | Table 3<br>Table 3 |
| Rubia et al., 2003 | 20 | TAL | stop signal task | control | general_go | CE | inhibition | Table 1 |
| Sharp et al., 2010 | 26 | MNI | stop signal task | control_continue | correct_continue | CE | inhibition | SM Table 4 |
| Sosic-Vasic et al., 2012 | 27 | MNI | go/no-go | correct | correct_stop | CE | inhibition | Table 3 |

| | | | | | | | | |
|---|---|---|---|---|---|---|---|---|
| Sozda et al., 2011 | 12 | TAL | stroop | control | correct_con_inc | GE | class. interference | Table 6 |
| Steele et al., 2014 | 102 | MNI | go/no-go | control,<br>correct | correct_go,<br>correct_stop | CE<br>CE | inhibition | Table 1,<br>Table 3 |
| Tang et al., 2006 | 18 | MNI | stroop | control,<br>control | correct_con_inc_neu,<br>correct_con_inc_neu | GE<br>GE | class. interference | Table 3,<br>Table 3 |
| Ullsperger and von Cramon, 2001 | 9 | TAL | flanker | correct | correct_inc | CE | class. interference | Table 3 |
| Wessel et al., 2012 | 21 | MNI | flanker | other | correct_standard | GE | class. interference | Table 1 |
| Wittfoth et al., 2008 | 15 | TAL | Simon | correct,<br>control | correct_inc,<br>correct_con | CE<br>GE | class. interference | Table 1,<br>Table 1 |
| Wittfoth et al., 2009 | 14 | TAL | simon | control | correct_con_inc_neu | GE | class. interference | Table 2 |
| Xu et al., 2017 | 21 | TAL | stop signal task | control | correct_go | CE | inhibition | SM Table 2 |
| Zandbelt et al., 2011 | 22 | MNI | stop signal task | correct | correct_stop | CE | inhibition | SM Table 8 |

Table S1: Overview of studies and respective experiments included in the meta-analysis on error processing. *Contrasted against* specifies against which control condition errors were contrasted (e.g. CE trials versus correct_go trials). Abbreviations: CE: Commission errors, GE: general errors, i.e. when not only commission errors but also omission errors were included in the regressor of interest. Different experiments from one study were pooled to account for sample-specific effects. * Studies using FSL or SPM and reporting TAL coordinates without stating that a transformation into TAL space was performed and hence treated as MNI space in the present analysis

| **Paper** | **n** | **MNI/TAL** | **name of task** | **contrasted against** | **type of task** | **Source of coordinates** |
|---|---|---|---|---|---|---|
| Aarts et al., 2008 | 12 | TAL | Stroop | correct_con | class. interference | Table 2 |
| Aron and Poldrack, 2006 | 13 | MNI | stop signal task | correct_go | inhibition | SM Table 2 |
| Basten et al., 2011 | 46 | MNI | Stroop | correct_con | class. interference | Table 3 |
| Becker et al., 2008 | 17 | TAL | Stroop | correct_con | class. interference | Table 2 |
| Berkman et al., 2014 | 60 | MNI | stop signal task | correct_go | inhibition | Table 2 |

| | | | | | | |
|---|---|---|---|---|---|---|
| Berron et al., 2015 | 24 | MNI | flanker | correct_con | class. interference | Table 2 |
| Bobb et al., 2012 | 13 | MNI | stop signal task | correct_go | inhibition | Table 2 |
| Boecker et al., 2011 | 15 | MNI* | stop signal task | correct_go | inhibition | SM Table 1 |
| Boehler et al., 2010 | 15 | MNI | stop signal task | go | inhibition | Table 3 |
| Bunge et al., 2002 | 16 | MNI* | flanker with NoGo component | correct_neu | class. interference | Table 1 |
| | | | flanker with NoGo component | correct_neu | inhibition | Table 2 |
| Cai and Leung, 2009 | 12 | MNI | stop signal task | correct_go (color task) | inhibition | Table 1 |
| | | | stop signal task | correct_go (orientation task) | inhibition | Table 1 |
| Cai and Leung, 2011 | 23 | MNI | stop signal task | go | inhibition | SM Table 1 |
| Cai et al. 2014 | 19 | MNI | stop signal task | correct_go | inhibition | Table 2 |
| Chen et al., 2015 | 25 | MNI | go/no-go | go | inhibition | Table 2 |
| Chen et al., 2018_1 | 36 | MNI | Stroop | correct_con | class. interference | Table 2 |
| Chen et al., 2018_2 | 35 | MNI | flanker | correct_con | class. interference | Table 2 |
| Chevrier et al., 2007 | 14 | TAL | stop signal task | go | inhibition | Table 1 |
| Chikara et al., 2018 | 20 | MNI | stop signal task | correct_go | inhibition | Table 3A |
| Chikazoe et al., 2007 | 24 | MNI | antisaccade | correct_control | class. interference | Table 1 |
| Chikazoe et al., 2009a | 25 | MNI | go/no-go | correct_frequent_go | inhibition | Table 1 |
| | | | go/no-go | correct_infrequent_go | inhibition | Table 2 |
| Chikazoe et al., 2009b | 22 | MNI | stop signal task | correct_uncertain_go | inhibition | Table 2 |

| | | | | | | |
|---|---|---|---|---|---|---|
| Christensen et al., 2011 | 26 | TAL | Stroop_Voice task | correct_neu (voice task) | class. interference | Table 2 |
| | | | Stroop_semantic task | correct_neu (semantic task) | class. interference | Table 2 |
| Chuang et al., 2014 | 60 | MNI | flanker | correct_con (high demand) | class. interference | Table 3 |
| | | | flanker | correct_con (low demand) | class. interference | Table 3 |
| Cieslik et al., 2010 | 24 | MNI | SRC | correct_con | class. interference | Text |
| Coderre et al., 2013 | 14 | MNI | Stroop | correct_con | class. interference | Table 1 |
| | | | Stroop | correct_neu | class. interference | Table 1 |
| Cojan et al., 2015 | 32 | MNI | flanker | correct_con | class. interference | Table 2 |
| Congdon et al., 2014 | 62 | MNI | stop signal task | correct_go | inhibition | Table 2 |
| Coxon et al., 2016_1 | 20 | MNI | stop signal task | correct_go | inhibition | Table 2 (young) |
| Coxon et al., 2016_2 | 20 | MNI | stop signal task | correct_go | inhibition | Table 2 (old) |
| Czapla et al., 2017 | 21 | MNI | go/no-go | correct_go | inhibition | SM Table 4 |
| | | | go/no-go | correct_go | inhibition | SM Table 5 |
| Daamen et al., 2015 | 100 | MNI | flanker | correct_con | class. interference | SM Table 3 |
| Dambacher et al., 2015 | 15 | TAL | go/no-go | correct_go | inhibition | Table 1 |
| Dichter et al., 2007 | 15 | MNI | flanker | correct_con (flanker) | class. interference | Table 2 |
| | | | flanker | correct_con (flanker_gaze) | class. interference | Table 2 |
| Fassbender et al., 2004 | 18 | TAL | go/no-go | tonic_activation | inhibition | Table 2 |
| Fauth-Bühler et al., 2012 | 18 | MNI | stop signal task | go/active baseline | inhibition | from authors |
| Fedota et al., 2014 | 16 | MNI | go/no-go | correct_go | inhibition | Table 1 |

| | | | | | | |
|---|---|---|---|---|---|---|
| Frühholz et al., 2011 | 24 | MNI | flanker color with Simon | correct_con | class. interference | Table 3d |
| Fuentes-Claramonte et al., 2016 | 57 | MNI | go/no-go | correct_frequent_go | inhibition | Table 2 |
| | | | go/no-go | correct_infrequent_go | inhibition | Table 2 |
| Ganos et al., 2014 | 15 | MNI | stop signal task | go | inhibition | SM Table 2 |
| Garavan et al., 1999 | 14 | TAL | go/no-go | go/active baseline | inhibition | Table 1 |
| Garavan et al., 2002 | 14 | TAL | go/no-go | go/active baseline | inhibition | Table 1 |
| Garavan et al., 2003 | 16 | TAL | go/no-go | go/active baseline | inhibition | Table 1 |
| Geng et al., 2009 | 16 | MNI | go/no-go | correct_go | inhibition | Table 3 |
| Ghahremani et al., 2012 | 18 | MNI | stop signal task | correct_go | inhibition | Table 3 |
| Grandjean et al., 2012 | 25 | MNI | Stroop | correct_neu | class. interference | Table 3a |
| Harle et al., 2016 | 34 | TAL | stop signal task | correct_go | inhibition | SM Table 2 |
| Hart et al., 2010 | 14 | MNI* | Stroop | correct_con | class. interference | Table 2 |
| Hester et al., 2004a | 15 | TAL | go/no-go | active baseline | inhibition | Table 3 |
| Hester et al., 2004b | 15 | TAL | go/no-go | active baseline | inhibition | Table 1 |
| Hinault et al., 2019 | 22 | MNI | Stroop | correct_con | class. interference | Table 2 |
| Holmes et al., 2010 | 33 | MNI | flanker | correct_con | class. interference | Table 3 |
| Holtmann et al., 2013 | 24 | MNI | flanker | correct_con | class. interference | Table 6 |
| Hough et al., 2016 | 22 | MNI | Stroop | correct_con | class. interference | SM Table |
| | | | go/no-go | correct_go | inhibition | SM Table |
| Hsu et al., 2017 | 20 | MNI | go/no-go | go | inhibition | Table 2 |
| Huang et al., 2012 | 30 | MNI | Stroop | correct_con | class. interference | Table 3 |
| Hughes et al., 2012 | 10 | MNI | stop signal task | active baseline | inhibition | Table 4 |

| | | | | | | |
|---|---|---|---|---|---|---|
| Hughes et al., 2013 | 15 | MNI | stop signal task | active baseline | inhibition | Table 3 |
| Iannaccone et al., 2015 | 15 | MNI | flanker | correct_con | class. interference | Table 2 |
| Ivanov et al., 2012 | 16 | MNI | flanker | correct_con | class. interference | Table 4 |
| Jahfari et al., 2011 | 20 | MNI | stop signal task | go | inhibition | Table 4 |
| Jahfari et al., 2012 | 16 | MNI | stop signal task | go | inhibition | Table 5 |
| Jahfari et al., 2015 | 23 | MNI | stop signal task | go | inhibition | Table 3 |
| Jaspar et al., 2014 | 45 | MNI | Stroop | correct_neu | class. interference | SM Table 2 |
| Jiang and Egner, 2014 | 21 | MNI | Stroop | correct_con | class. interference | Table 1 |
| | | | Simon | correct_con | class. interference | Table 1 |
| Kaladjian et al., 2007 | 21 | TAL | go/no-go | correct_go | inhibition | Table 3 |
| Kaladjian et al., 2009a | 10 | TAL | go/no-go | correct_go (t1) | inhibition | Table 3 |
| | | | go/no-go | correct_go (t2) | inhibition | Table 3 |
| Kaladjian et al., 2009b | 20 | TAL | go/no-go | correct_go | inhibition | Table 3 |
| Kelly et al., 2004 | 15 | TAL | go/no-go | active baseline | inhibition | Table1 |
| Kenner et al., 2010 | 24 | MNI | stop signal task | correct_go | inhibition | SM Table 3 |
| Kerns et al., 2005 | 13 | TAL | Stroop | correct_con | class. interference | Table 1 |
| Kiehl et al., 2000 | 14 | MNI | go/no-go | active baseline | inhibition | Table 1 |
| Kim et al., 2011 | 13 | MNI | Stroop | correct_neu | class. interference | Table 1 |
| Kim et al., 2012 | 16 | MNI | Stroop | correct_con | class. interference | Table 2 |
| Kim et al., 2014 | 18 | MNI | Stroop | correct_con | class. interference | Table 1 |
| King et al., 2012 | 25 | MNI | flanker | correct_con | class. interference | Table 1 |
| Ko et al., 2014 | 23 | MNI | go/no-go | go | inhibition | Table 2 |
| Ko et al., 2016 | 32 | MNI | stop signal task | correct_go (simple symbol) | inhibition | Table 1A |
| | | | stop signal task | correct_go (batterfield symbol) | inhibition | Table 1B |
| Köhler et al., 2018 | 33 | MNI | go/no-go | correct_go | inhibition | SM Table 1 |

| | | | | | | |
|---|---|---|---|---|---|---|
| Korsch et al., 2014_1 | 20 | MNI | flanker with SRC | correct_con | class. interference | Table 2 (young) |
| Korsch et al., 2014_2 | 19 | MNI | flanker with SRC | correct_con | class. interference | Table 2 (old) |
| Krönke et al., 2018 | 139 | MNI | Stroop | correct_con | class. interference | Table 7 |
| Lavallee et al., 2014 | 21 | MNI | stop signal task | go | inhibition | Table 2 |
| Lee et al., 2008 | 14 | MNI | Simon | correct_con | class. interference | Table 3 |
| Lenartowicz et al. 2011 | 23 | MNI | stop signal task | correct_go (1) | inhibition | Table 2 |
| | | | stop signal task | correct_go (2) | inhibition | Table 2 |
| Lesh et al., 2013 | 54 | MNI | Stroop | correct_con | class. interference | Table 2 |
| Liddle et al. , 2001 | 16 | MNI | go/no-go | correct_go | inhibition | Table 3 |
| Lorenz et al., 2015 | 38 | MNI | stop signal task | go | inhibition | SM Table 1 |
| Luethi et al., 2016 | 88 | MNI | Stroop | correct_con | class. interference | SM Table 2 |
| Luks et al., 2007 | 11 | TAL | flanker | correct_con | class. interference | Table 3 |
| Lütcke et al., 2008 | 12 | MNI | flanker | correct_con | class. interference | Table 2 |
| Manard et al., 2017 | 40 | MNI | Stroop | correct_neu | class. interference | SM Table 1A |
| Marco-Pallarés et al., 2008 | 10 | MNI | stop signal task | correct_go | inhibition | Table 1 |
| Mazzola-Pomietto et al., 2009 | 16 | TAL | go/no-go | correct_go | inhibition | Table 3 |
| Meffert et al., 2016 | 22 | TAL | go/no-go | correct_go | inhibition | Table 1 |
| Mohammadi et al., 2015 | 17 | MNI | stop signal task | go | inhibition | SM Table 2 |
| Montojo et al., 2013 | 30 | MNI | stop signal task | correct_go | inhibition | Table 2 |
| Nakata et al., 2008 | 15 | TAL | go/no-go | correct_go | inhibition | Table 5 |
| O'Connor et al., 2012 | 18 | MNI | go/no-go | active baseline | inhibition | Table 1 |
| Overbeek et al., 2019 | 21 | MNI | Stroop | correct_con | class. interference | SM Table 1 |
| Page et al., 2009 | 11 | TAL | go/no-go | correct_go | class. interference | Appendix A Table |

| | | | | | | |
|---|---|---|---|---|---|---|
| | | | Simon | correct_con | class. interference | Appendix A Table |
| Paschke et al., 2015 | 115 | MNI | flanker | correct_con | class. interference | Table 2 |
| Piai et al., 2013 | 23 | MNI | Stroop | correct_con | class. interference | Table 6 |
| | | | Stroop | correct_neu | class. interference | Table 6 |
| Pompei et al., 2011 | 48 | TAL | Stroop | correct_neu | class. interference | SM Table |
| Prakash et al., 2009_1 | 25 | MNI | Stroop | correct_neu | class. interference | Table 2a (old) |
| Prakash et al., 2009_2 | 25 | MNI | Stroop | correct_neu | class. interference | Table 2b (young) |
| Purmann & Pollmann, 2015 | 18 | MNI | Stroop | correct_con | class. interference | Table 2 |
| Rae et al., 2014 | 17 | MNI | stop signal task | correct_go (specified) | inhibition | SM Table 2 |
| | | | stop signal task | correct _go (select) | inhibition | SM Table 2 |
| Ramautar et al., 2006 | 16 | MNI | stop signal task | go | inhibition | Table 1 |
| Ramm et al., 2020 | 20 | MNI | Stroop | correct_con | class. interference | SM Table 2 |
| Rodriguez-Pujadas et al., 2014 | 33 | MNI* | stop signal task | correct_go | inhibition | Table 2 |
| Rothmayr et al., 2011 | 12 | MNI | go/no-go | correct_go | inhibition | Table 2 |
| Rubia et al., 2006 | 21 | TAL | go/no-go | correct_go | inhibition | Table 2 |
| | | | Simon | correct_con | class. interference | Table 2 |
| Schel et al., 2014 | 24 | MNI | stop signal task | correct_go | inhibition | Table 4 |
| Schmidt et al., 2012 | 31 | MNI | Stroop | correct_con | class. interference | from authors |
| Sebastian et al., 2012 | 24 | MNI | go/no-go | active baseline | inhibition | Table 5 |
| | | | stop signal task | correct_go | inhibition | Table 5 |
| | | | Simon | correct_con | class. interference | Table 5 |
| Sebastian et al., 2013a | 48 | MNI | Simon | correct_con | class. interference | SM Table 1 |
| | | | go/no-go | active baseline | inhibition | SM Table 1 |

| | | | | | | |
|---|---|---|---|---|---|---|
| | | | stop signal task | correct_go | inhibition | SM Table 1 |
| Sebastian et al., 2013b_1 | 24 | MNI | Simon | correct_con | class. interference | Table 3 |
| | | | go/no-go | active baseline | inhibition | Table 3 |
| | | | stop signal task | correct_go | inhibition | Table 3 |
| Sebastian et al., 2013b_2 | 21 | MNI | Simon_hybrid response interference | correct_congruent go | class. interference | Table 2 |
| | | | go/no-go_hybrid response interference | correct_congruent go | inhibition | Table 2 |
| | | | stop signal task_hybrid response interference | correct_congruent go | inhibition | Table 2 |
| Sebastian et al., 2017 | 80 | MNI | stop signal task | correct_go | inhibition | from authors |
| Sharp et al., 2010 | 26 | MNI | stop signal task | correct_go | inhibition | SM Table 1 |
| Shin & Kim, 2015 | 43 | MNI | Stroop | correct_con | class. interference | Table 1 |
| Steele et al. 2014 | 102 | MNI | go/no-go | correct_go | inhibition | Table 2 |
| Swann et al. 2012 | 16 | MNI | stop signal task | correct_go | inhibition | SM Table 2 |
| Tabu et al., 2011 | 13 | MNI | stop signal task | correct_go | inhibition | text, page 280 |
| Tabu et al., 2012 | 13 | MNI | stop signal task | correct_go | inhibition | SM Table 1 (hand task) |
| | | | stop signal task | correct_go | inhibition | SM Table 1 (foot task) |
| Trautwein et al., 2016 | 282 | MNI | flanker | correct_con | class. interference | Table 1 |
| Ullsperger et von Cramon, 2001 | 9 | TAL | flanker | correct_con | class. interference | Table 2 |
| Van der Meer et al., 2013 | 19 | MNI | stop signal task | correct_go | inhibition | SM Table 3A |
| Van Eijk et al., 2015_1 | 18 | MNI | go/no-go | correct_go | inhibition | Table 4 |
| | | | stop signal task | correct_go | inhibition | Table 4 |

| | | | | | | |
|---|---|---|---|---|---|---|
| | | | Simon | correct_con | class. interference | Table 4 |
| van Eijk et al., 2015_2 | 25 | MNI | Simon_hybrid response interference | correct_congruent go | class. interference | Table 5 |
| | | | go/no-go_hybrid response interference | correct_congruent go | inhibition | Table 5 |
| | | | stop signal task_hybrid response interference | correct_congruent go | inhibition | Table 5 |
| Verstynen et al., 2014 | 28 | MNI | Stroop | correct_neu | class. interference | Table 1 |
| Wallentin et al., 2015 | 49 | MNI | Stroop | correct_con | class. interference | from authors |
| Walther et al., 2010 | 17 | MNI | go/no-go | correct_go (audi AND visual) | inhibition | Table 1 |
| Wang et al., 2014 | 20 | MNI | flanker | correct_con | class. interference | Table 4 |
| Wilbertz et al., 2014 | 49 | MNI | stop signal task | go | inhibition | SM Table 2 |
| Wittfoth et al., 2008 | 15 | TAL | Simon | correct_con | class. interference | Table 1 |
| Xu et al., 2015 | 18 | MNI | stop signal task | go | inhibition | Table 2 |
| Xu et al., 2017 | 21 | MNI* | stop signal task | correct_go | inhibition | SM Table 1 |
| Xue et al., 2008 | 15 | MNI | stop signal task | correct_go | inhibition | SM Table 1 (manual) |
| | | | stop signal task | correct_go | inhibition | SM Table 1 (letter naming) |
| Ye and Zhou., 2009 | 19 | MNI | flanker | correct_con | class. interference | Table 3 |
| | | | Stroop | correct_con | class. interference | Table 3 |
| Zandbelt and Fink, 2010 | 24 | MNI | stop signal task | go | inhibition | SM Table 3B |
| Zandbelt et al. 2011 | 22 | MNI | stop signal task | go | inhibition | SM Table 11 |
| Zheng et al., 2008 | 18 | TAL | go/no-go | go | inhibition | Table 1 |

| | | | | | | |
|---|---|---|---|---|---|---|
| | | | stop signal task | go | inhibition | Table 1 |
| Zurawska vel Grajewska et al., 2011 | 18 | MNI | flanker | correct_con | class. interference | Table 2b |

SM Table 2: Overview of studies included in the meta-analysis on successful interference control. *Contrasted against* specifies against which control condition successful interference responses were contrasted (e.g., correct stop trials versus correct go trials). Different experiments from one study were pooled to account for sample-specific effects. If the same study reported two independent participant samples, the respective experiments were treated as independent experiments (e.g. Chen et al., 2018_1 and Chen et al., 2018_2). * Studies using FSL or SPM and reporting TAL coordinates without stating that a transformation into TAL space was performed and hence treated as MNI space in the present analysis
